# WIND1 controls cell fate transition through histone acetylation and deacetylation during somatic embryogenesis

**DOI:** 10.1101/2025.08.07.669221

**Authors:** Akira Iwase, Arika Takebayashi, Fu-Yu Hung, Ayako Kawamura, Yetkin Çaka Ince, Yasuhiro Kadota, Soichi Inagaki, Takamasa Suzuki, Ken Shirasu, Keiko Sugimoto

## Abstract

Regeneration involves large-scale transcriptional reprogramming to drive cell identity transitions. These transcriptional changes are tightly coupled with chromatin remodelling but molecular mechanisms that coordinate these changes remain unclear. Here we show that WOUND INDUCED DEDIFFERENTIATION 1 (WIND1) transcription factor promotes somatic embryogenesis by repressing pre-existing cell fate and activating new cell identity programmes. WIND1 interacts with histone deacetylase HISTONE DEACETYLASE 9 (HDA9) and histone acetyltransferase complex component HOMOLOG OF YEAST ADA1 2a (ADA2a) *via* conserved N-terminal domain. These interactions enable WIND1 to mediate both H3K27 deacetylation and acetylation at distinct target loci, leading to repression of shoot identity genes such as *AINTEGUMENTA* (*ANT*) and activation of embryogenesis regulators including *LEAFY COTYLEDON 2* (*LEC2*). Our findings identify WIND1 as a bifunctional chromatin regulator that integrates opposing histone acetylation dynamics to coordinate transcriptional reprogramming. This mechanism provides a molecular framework for how a transcription factor directs complex cell fate transitions during regeneration

## INTRODUCTION

Regeneration involves dynamic changes in cell fate, requiring the erasure of original cellular identity and acquisition of new developmental programmes (Birnbaum and Alvarado, 2008). These transitions require coordinated changes at the molecular level, including signalling cascades, chromatin remodelling, and transcriptional control. Among these layers, the regulation of gene expression has emerged as a central mechanism that integrates external cues with intrinsic regulatory states to direct cell fate transitions. In animals, induced pluripotency through reprogramming factors like Octamer-binding transcription factor 4 (Oct4), SRY-box transcription factor 2 (Sox2), Krüppel-like factor 4 (Klf4), and c-Myc demonstrates differentiated cells can regain developmental potential (Takahashi and Yamanaka, 2006). Similar principles underlie regeneration in organisms such as salamanders and zebrafish, where mature cells re-enter proliferative states and acquire developmental plasticity before adopting new fates (Tanaka and Reddien, 2011; Jopling et al., 2011). Central to these transitions are transcription factors that both activate new gene expression programmes and suppress prior identities (Chronis et al., 2017). In plants, overexpression of certain transcription factors can bypass developmental restrictions and trigger pluripotency or totipotency states. LEAFY COTYLEDON 1 (LEC1), WUSCHEL (WUS), and BABY BOOM (BBM), for example, can induce somatic embryogenesis or shoot regeneration when ectopically expressed (Lotan et al., 1998; Boutilier et al., 2002; Zuo et al., 2002).

Wounding serves as a common physiological trigger for initiating such transitions by inducing dedifferentiation, a process in which differentiated cells revert to a more plastic, stem cell-like state (Tanaka and Reddien, 2011; Ikeuchi et al., 2013; Ikeuchi et al., 2016). We previously identified an AP2/ERF transcription factor WOUND INDUCED DEDIFFERENTIATION 1 (WIND1) as a key regulator of wound-induced regeneration, promoting callus formation, shoot regeneration, and vascular reconnection in *Arabidopsis thaliana* (Arabidopsis) (Iwase et al., 2011; Iwase et al., 2017; Iwase et al., 2021). Recent work in tomato revealed that peptide-receptor signalling relays mechanical cues from wounds to *WIND1* expression, establishing a local signalling module that links mechanical stress to transcriptional reprogramming (Yang et al., 2024).

In addition to the regulation by transcription factors, epigenetic regulation, particularly histone modifications, plays pivotal roles in cell reprogramming(Chronis et al., 2017; Wang et al., 2020; Chen et al., 2023; Liu et al., 2023). Among these, histone acetylation and deacetylation dynamically modulate chromatin accessibility, thereby influencing gene activity (Bannister and Kouzarides, 2011). Acetylation of histone lysine residues, mediated by histone acetyltransferases (HATs), relaxes chromatin structure and promotes gene activation, whereas deacetylation by histone deacetylases (HDACs) results in chromatin condensation and transcriptional repression. These modifications are dynamic and responsive to developmental and environmental cues. In plants, histone acetylation has been implicated in stress-induced reprogramming and developmental plasticity, including organ regeneration (Ikeuchi et al., 2019; Oberkofler et al., 2021; Sakamoto et al., 2022; Li et al., 2024). Increasing evidence suggests that transcription factors can recruit HATs and HDACs to specific genomic loci to control regeneration-associated gene expression (Zhou et al., 2017; Lee et al., 2024), suggesting that transcription factors actively engage chromatin regulators to mediate cell fate transitions. Somatic embryogenesis, in which embryos arise from somatic cells, offers a powerful model to study the mechanisms of plant cell fate reprogramming. Somatic embryos can be induced from immature zygotic embryos by exogenous phytohormones such as auxins (Gaj, 2001). This regenerative potential is, however, sharply reduced after germination due to chromatin-mediated transcriptional repression of key regulators such as *LEC2* (Stone et al., 2001; Wang et al., 2020). Interestingly, application of stressors such as osmotic pressure and heat shock can restore somatic embryogenesis competence by reactivating these genes (Ikeda-Iwai et al., 2003; Kadokura et al., 2018), although the mechanisms that overcome these post-embryonic barriers to initiate embryogenesis remain unclear.

While the acquisition of new cell identities has been a major focus of regeneration studies, dedifferentiation is equally essential. Dedifferentiation constitutes a critical early phase of regeneration across systems, from salamander limb regrowth to callus formation in Arabidopsis (Tanaka and Reddien, 2011; Ikeuchi et al., 2013; Sugiyama, 2015). Little is known, however, about how dedifferentiation is mechanistically regulated or how it is coordinated with the induction of new fates. In this study we address this gap by using a WIND1-inducible culture system. Our data show that WIND1 is both necessary and sufficient to induce somatic embryogenesis in Arabidopsis. WIND1 acts through a dual chromatin-based mechanism since it interacts with HISTONE DEACETYLASE 9 (HDA9) to repress genes associated with shoot identity and with ADA2a-HAG1 histone acetyltransferase complex to activate embryonic regulators. Our findings reveal a transcription factor–chromatin module that integrates environmental signals with histone acetylation dynamics to coordinate cell fate transitions during regeneration.

## RESULTS

### WIND1 is both sufficient and necessary for somatic embryogenesis

As previously reported (Ikeuchi et al., 2013), ectopic expression of *WIND1* in *XVE:WIND1* seedlings treated with 17β-estradiol (17ED) induces the formation of embryo-like tissues around the shoot apical meristem (Figures 1A and 1B). These tissues exhibit characteristic features of embryos, including cotyledons, hypocotyls and juvenile root-like structures, and they subsequently develop into complete plantlets after transfer to 17ED-free conditions (Figure 1A). Moreover, these embryo-like tissues can be stained with Sudan Red, a dye that marks lipid accumulation in the embryonic tissues (Figure 1B) (Bouyer et al., 2011), confirming that increased *WIND1* expression is sufficient to induce embryo formation from somatic tissues.

**Figure 1.**
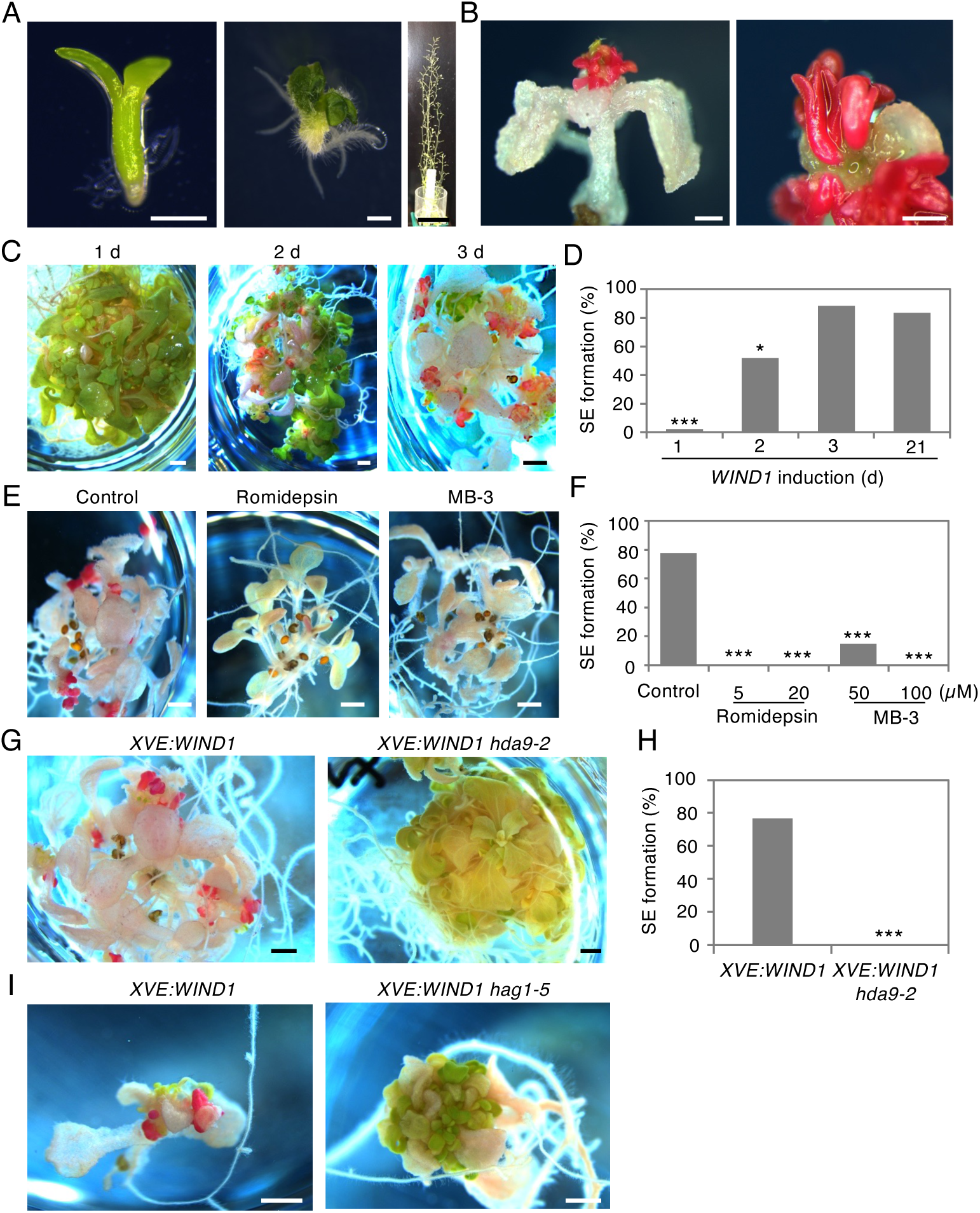
HDA9 and HAG1 are required for WIND1-induced somatic embryogenesis. (A) An embryo-like tissue isolated from 17β-estradiol (17ED)-treated *XVE:WIND1* seedlings (left), true leaves and roots formed after transfer to 17ED-free MS medium (middle) and fertile plants grown on soil (right). Scale bars, 500 μm (left, middle), 5 cm (right). (B) Sudan Red-stained somatic embryos on *XVE:WIND1* seedlings treated with 17ED for 21 days (left) and magnified image (right). Scale bars, 500 μm. (C and D) Sudan Red-stained somatic embryos on *XVE:WIND1* seedlings treated with 17ED for 1, 2, 3, or 21 days (d) (C). Quantitative data of somatic embryo (SE) formation shown as ratio (%) of seedlings that formed SE among all tested seedlings (D). Scale bars, 1mm. Statistical significance was determined by proportion test (n = 50 for 1 d and 2 d, 42 for 3 d, 54 for 21 d treatment, \**p* < 0.005, \*\*\**p* < 3.75e-16). (E and F) Sudan Red-stained somatic embryos on 17ED-treated *XVE:WIND1* seedlings in the presence of 5 µM romidepsin, a class I histone deacetylase inhibitor, or 100 µM MB-3, a histone acetyltransferase inhibitor (E). Quantitative data of SE formation (F). Scale bars, 1mm. Statistical significance was determined by proportion test (n = 71 for control, 64 for romidepsin, 67 for MB-3, \*\*\**p* < 1.92e-11). (G and H) Sudan Red-stained somatic embryos on 17ED-treated *XVE:WIND1* and *XVE:WIND1 hda9* seedlings (G). Quantitative data of SE formation (H). Scale bars, 1mm. Statistical significance was determined by proportion test (n = 69 for *XVE:WIND1* and 67 for *XVE:WIND1 hda9-2*. \*\*\**p* < 2.2e-16). (I) Sudan Red-stained somatic embryos on 17ED-treated *XVE:WIND1* and *XVE:WIND1 hag1-5* seedlings. Scale bars, 1 mm. See also Supplemental Figures 1-3.

To further define the temporal requirement for WIND1 activity, we examined the duration of *WIND1* induction necessary to trigger somatic embryo formation in this system. A one-day induction was insufficient to induce somatic embryos whereas 2 or more days of induction successfully triggered their formation (Figures 1C and 1D). Notably, *XVE:WIND1* seedlings continued to form true leaves around the shoot apical meristem after 1 or 2 days of induction, indicating that at least 3 days of WIND1 activation is required to fully suppress shoot identity and redirect the developmental programme into embryogenesis.

Previous work by Wang et al. (2020) showed that *WIND1* expression increases at the early stages of hormone-induced somatic embryogenesis from immature embryos (Wang et al., 2020), implying a functional role of WIND1 in this process. To test this idea, we performed the hormone-induced somatic embryogenesis assay (Gaj, 2001; Wang et al., 2020), using immature embryos isolated from *WIND1-SRDX* plants expressing the dominant negative form of WIND1(Iwase et al., 2011). As shown in Supplemental Figures 1A-C, somatic embryo formation was significantly reduced in *WIND1-SRDX* plants compared to wild-type (WT) plants, establishing that WIND1 is not only sufficient but also necessary for the initiation of somatic embryogenesis.

### HDA9 and HAG1 are required for WIND1-induced somatic embryogenesis

To investigate whether WIND1 influences histone acetylation dynamics during somatic embryogenesis, we first examined the effects of chemical inhibitors targeting HDACs and HATs (Sako et al., 2015; Ueda et al., 2017; Rymen et al., 2019). As shown in Figures 1E, 1F, Supplemental Figures 2A and 2B, treatment with class I HDAC inhibitors such as romidepsin and Ky-2, as well as the HAT inhibitor γ-butyrolactone (MB-3), significantly suppressed somatic embryo formation in the WIND1-induction system. These results suggest that WIND1 promotes embryogenesis by modulation of both histone deacetylation and acetylation, likely affecting histone H3 marks.

**Figure 2.**
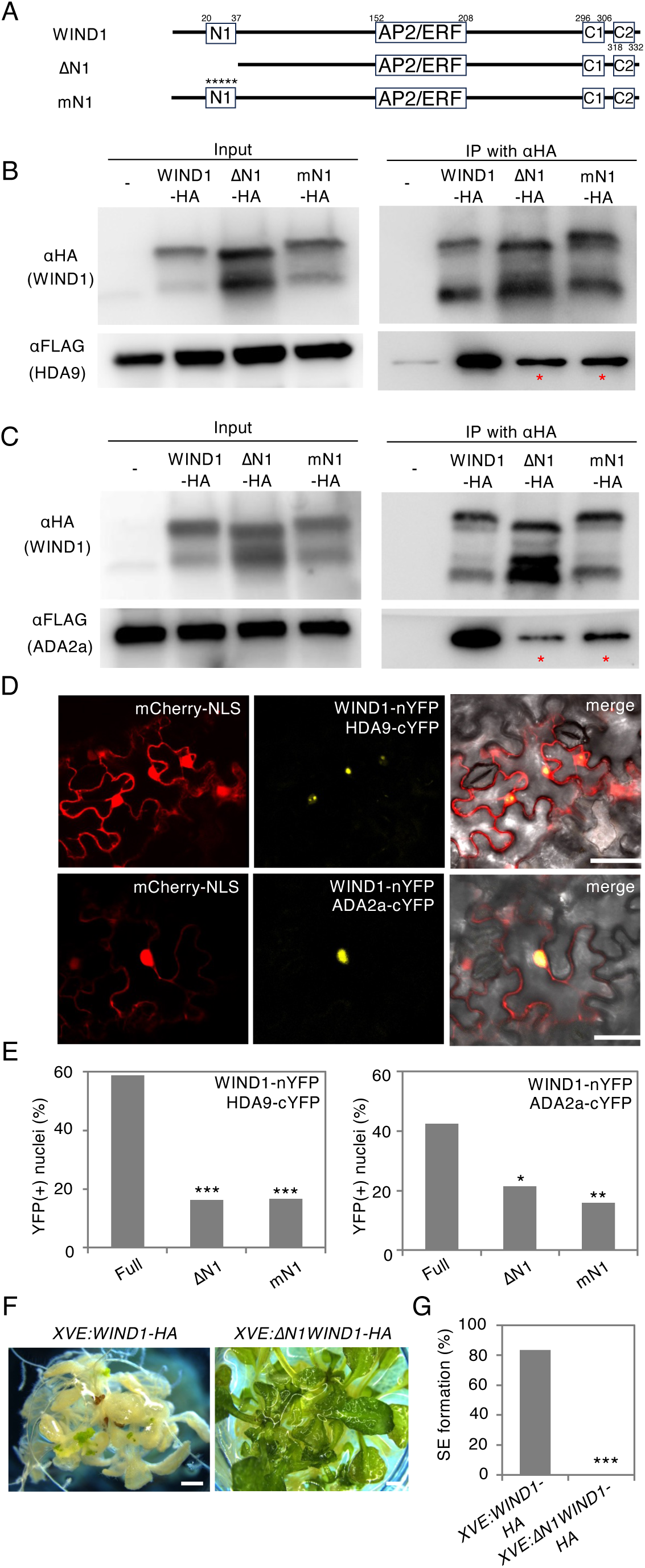
WIND1 directly interacts with HDA9 and ADA2a *in vivo*. (A) Schematic diagram of *WIND1* constructs. White boxes represent amino acid sequences conserved among WIND1 homologues (Iwase et al., 2013) and numbers above boxes show the order of amino acids. N1; conserved N-terminal domain 1, AP2/ERF; DNA binding domain, C1 and C2; conserved C-terminal domain 1 and 2. WIND1; WIND1 full-length, ΔN1; WIND1 lacking from 1^st^ to the 37^th^ amino acids, mN1; WIND1 with amino acid substitution within the N1 domain. Asterisks highlight the substitution of two leucines (L21A, L25A) and three serines (S34A, S36A, S37A) with alanines (A) within the N1 domain. (B and C) Immunoprecipitation assay showing in vivo interactions of WIND1 to HDA9 (B) and ADA2a (C). Soluble extracts (input) from agroinfiltrated *N. benthamiana* leaves co-expressing WIND1-HA or its variants (ΔN1-HA and mN1-HA) with HDA9-FLAG (B) or with ADA2a-FLAG (C) were subject to immunoprecipitation with αHA antibody, followed by immunoblotting with αHA and αFLAG antibodies. Asterisks highlight reduced levels of HDA9-FLAG and ADA2a-FLAG co-immunoprecipitated with ΔN1-HA or mN1-HA compared to full-length WIND1-HA. (D) BiFC assays in *N. benthamiana* epidermal cells showing the interaction between WIND1 and HDA9, or WIND1 and ADA2a within the nucleus. WIND1 fused with the N terminus of YFP (nYFP) and HAD9 or ADA2a fused with the C terminus of YFP (cYFP) were co-expressed into *N. benthamiana* leaves. The nucleus was marked by mCherry carrying a nuclear localization signal (NLS). Scale bars, 50µm. (E) Quantitative analysis of BiFC assays in (D). Statistical significance was determined by proportion test (n > 97 for each condition. \**p* < 0.005; \*\**p* < 2.2e-5; \*\*\**p* < 1.7e-9). (F and G) *XVE:WIND1-HA* and *XVE:ΔN1WIND1* seedlings treated with 17ED for 21 days, just before Sudan Red staining (F). Note that *XVE:ΔN1WIND1* seedlings do not display de-greening and suppression of leaf development. Quantitative data of SE formation (G). Scale bars, 1mm. Statistical significance was determined by proportion test (n=36 for *XVE:WIND1-HA*, 53 for *XVE:ΔN1WIND1*. \*\*\**p* < 2.96e-10). See also Supplemental Figures 4-6.

Given that the commitment for somatic embryogenesis occurs within 3 days of WIND1 induction (Figures 1C and 1D), we next tested when these chromatin-modifying activities are required. Interestingly, addition of romidepsin within the first day of *WIND1* activation was sufficient to nearly abolish somatic embryogenesis (Supplemental Figures 2C and 2E), indicating that HDAC activity is essential during the early phase of *WIND1* induction. Romidepsin-treated *XVE:WIND1* seedlings continued to develop true leaves (Supplemental Figures 2C), suggesting that HDAC activity contributes to the suppression of shoot identity following *WIND1* induction. In contrast, treatment with MB-3 for 1 or 2 days only partially inhibited somatic embryogenesis and 3-day exposure was required for complete suppression (Supplemental Figures 2D and 2F). These results suggest that HAT activity is required in multiple stages during embryogenic reprogramming.

The wound-induced callus formation assay is a quick and versatile system for identifying factors involved in stress-induced cell reprogramming (Iwase et al., 2017; Iwase et al., 2021). Using this assay we previously showed that *histone acetyltransferase of the gnat/myst superfamily 1/general control nonrepressed 5* (*hag1/gcn5*) and *hag3* mutants display severe defects in callus formation at wound sites (Supplemental Figures 3A and 3B)(Rymen et al., 2019). To identify class I HDACs involved in WIND1-mediated cell reprogramming, we screened for a collection of Arabidopsis HDAC mutants. Among them, two null alleles of HISTONE DEACETYLASE 9 (HDA9), *hda9-1* and *hda9-2*, displayed significantly reduced wound-induced callus formation (Supplemental Figures 3A and 3B), suggesting a potential function in reprogramming. To test the role of HDA9 in WIND1-induced somatic embryogenesis, we introduced the *hda9-2* mutation into the *XVE:WIND1* background by genetic crosses. Strikingly, the *hda9-2* mutation completely abolished somatic embryo formation upon *WIND1* induction (Figure 1G and 1H). We also attempted to generate *XVE:WIND1 hag1-5* lines but this proved challenging due to the severely reduced seed set of *hag1-5* mutants. Nevertheless, the few *XVE:WIND1 hag1-5* plants we obtained failed to form embryos and instead produced small true leaves, indicating a failure to acquire embryonic identity (Figure 1I). Together, these results provide genetic evidence that both HDA9 and HAG1 are essential for WIND1-induced somatic embryogenesis.

**Figure 3.**
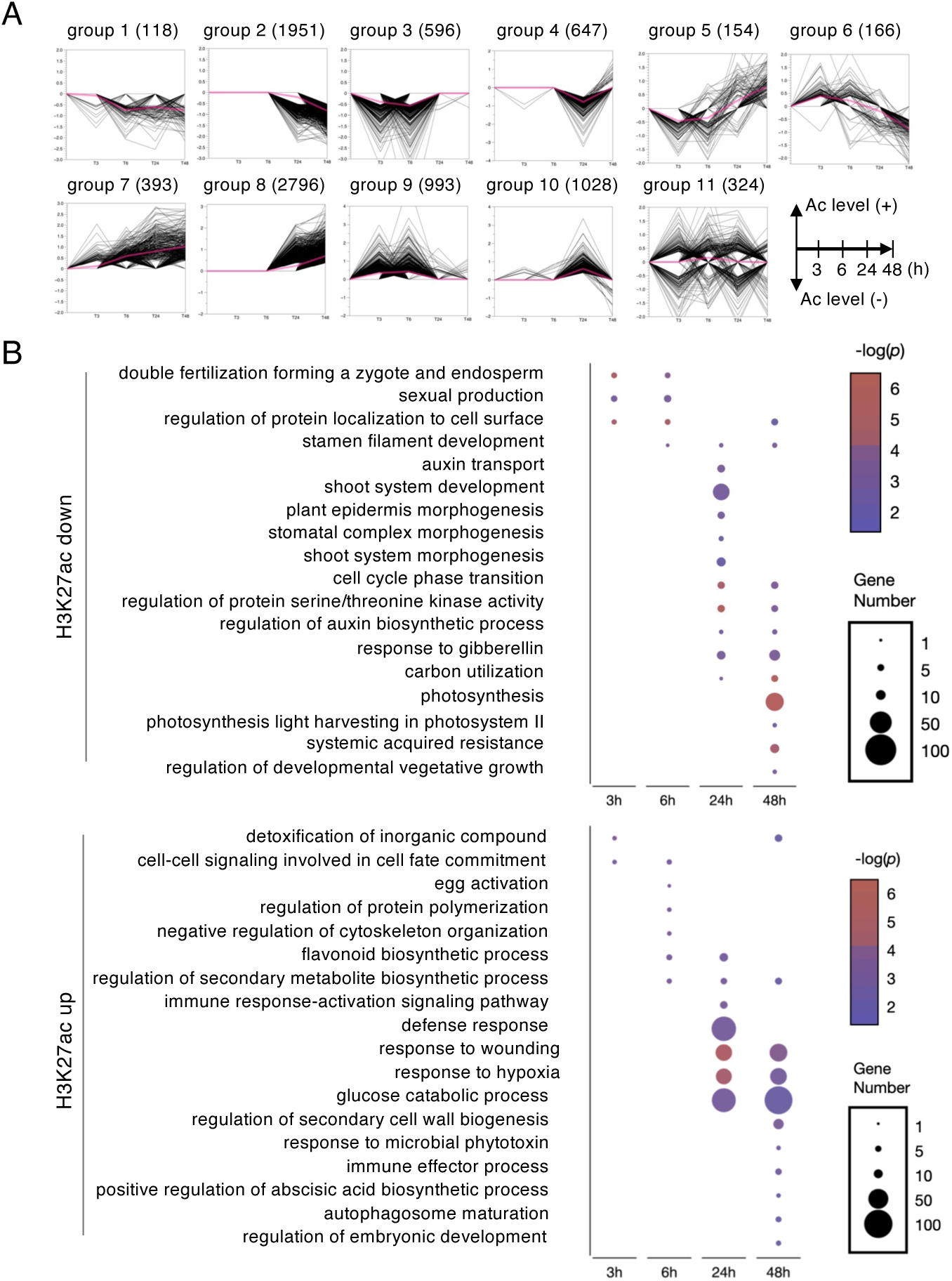
WIND1 induces histone deacetylation or acetylation at specific target loci. (A) Temporal dynamics of H3K27ac enrichment levels at 0, 3, 6, 24 and 48 h after WIND1 induction. Genes showing significant changes in H3K27ac level compared to H3 level (Student’s t-test, *p* < 0.05) were clustered into 11 groups. Number of genes included in each group is shown in brackets and red lines mark their average H3K27ac levels. Genes that show transient or constitutive reduction of H3K27ac levels in groups 1 to 4 and genes that show transient or constitutive increase of H3K27ac levels in groups 7 to 10 were further characterized in (B). (B) GO enrichment analysis of genes with reduced (Fold change <0.7, *p* <0.05) or elevated H3K27ac levels (Fold chang >1.3, *p* <0.05) following WIND1 induction, compared to the 0 h time point. Circle size and colour indicate the number of genes and statistical significance (*p*-value from Fisher’s exact test), respectively. See also Supplemental Tables 1 and 2.

### WIND1 interacts with HDA9 and ADA2 adaptor proteins that associate with HAG1

To explore how WIND1 modulates HDA9- and HAG1-mediated histone acetylation dynamics, we first examined whether it regulates the expression of these genes. Time-course microarray data on 17ED-treated *XVE:WIND1* seedlings, however, did not show significant changes in the transcript levels of these genes within 24 hour after *WIND1* induction (Iwase et al., 2021), suggesting that WIND1 does not act through transcriptional regulation for these histone acetylation regulators.

We therefore hypothesised that WIND1 might interact with these enzymes and recruit them to specific genomic loci to locally alter histone acetylation. To test this hypothesis, we conducted the yeast two-hybrid (Y2H) assays. Since full-length WIND1 exhibited autoactivation in the Y2H system, we used a truncated version that lacked the C-terminal C2 domain (ΔC2WIND1) as bait (Supplemental Figure 4A). As shown in Supplemental Figure 4B, we detected direct interactions between WIND1 and HDA9 in the yeast cells. In contrast, WIND1 did not directly interact with HAG1 but instead bound to HOMOLOG OF YEAST ADA2 2a (ADA2a) or ADA2b, adaptor proteins known to associate with HATs including HAG1 (Supplemental Figure 4B) (Mao et al., 2006; Anzola et al., 2010; Wu et al., 2023). To determine the domain of WIND1 responsible for the interaction with HDA9 and ADA2a, we generated 3 truncated versions of the WIND1 protein, each containing the N-terminal region, the DNA binding domain or the C-terminus region (Supplemental Figure 5A). Intriguingly, we found that both HDA9 and ADA2a interact specifically with the N-terminal region of WIND1 (Supplemental Figure 5B).

To validate these interactions *in planta*, we performed the co-immunoprecipitation (Co-IP) experiments using *Nicotiana benthamiana* transient expression system. Consistent with the Y2H results, WIND1 interacted with both HDA9 and ADA2a through its N-terminal region since the truncated form of WIND1 lacking the N-terminal 151 amino acid (ΔNWIND1) exhibited markedly reduced binding to both HDA9 and HAG1 (Supplemental Figures 5C-5E). Notably, this N-terminal region includes a short stretch of amino acids from glutamate at position 20 to serine at position 37 that is highly conserved among WIND1 orthologs across various plant species and referred to as the N1 domain (Iwase et al., 2013). Deletion of the N1 domain (ΔN1WIND1) or alanine substitution of two conserved leucines (L21 and L25) and three serines (S33, S35 and S37) (mN1WIND1) severely impaired binding to HDA9 and ADA2a (Figures 2A-C). These results indicate that WIND1 interacts with HDA9 and ADA2a specifically through its conserved N1 domain.

**Figure 4.**
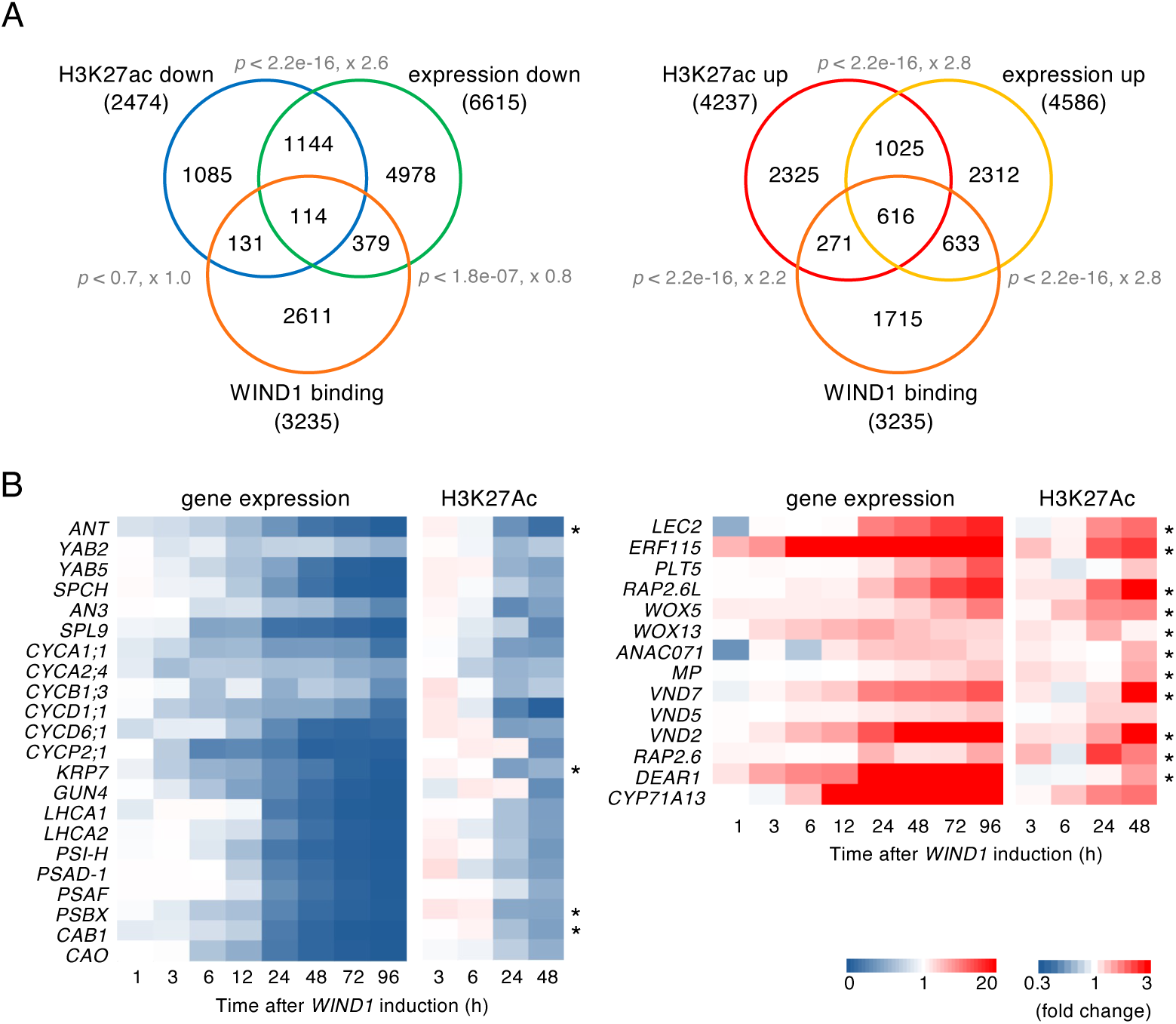
WIND1 directly regulates acetylation-coupled gene expression. (A) Venn diagrams showing the overlap among genes that show reduced H3K27ac, reduced gene expression and WIND1 binding (left) or genes that show increased H3K27ac, increased gene expression and WIND1 binding (right) after WIND1 induction in *XVE:WIND1* or *XVE:WIND1-HA* plants. The statistical significance of overlap between each pair of gene sets was assessed using Fisher’s exact test; corresponding *p*-values and odds ratios are shown. (B) Heatmap showing the temporal dynamics of gene expression at 1, 3, 6, 12, 24 and 48, 72 and 96 h after WIND1 induction as well as H3K27ac enrichment levels at 3, 6, 24 and 48 h after WIND1 induction. Values are shown as relative to the 0 h time point. Asterisks indicate direct WIND1 binding based on ChIP-seq data. See also Supplemental Figure 7 and Supplemental Tables 1-6.

**Figure 5.**
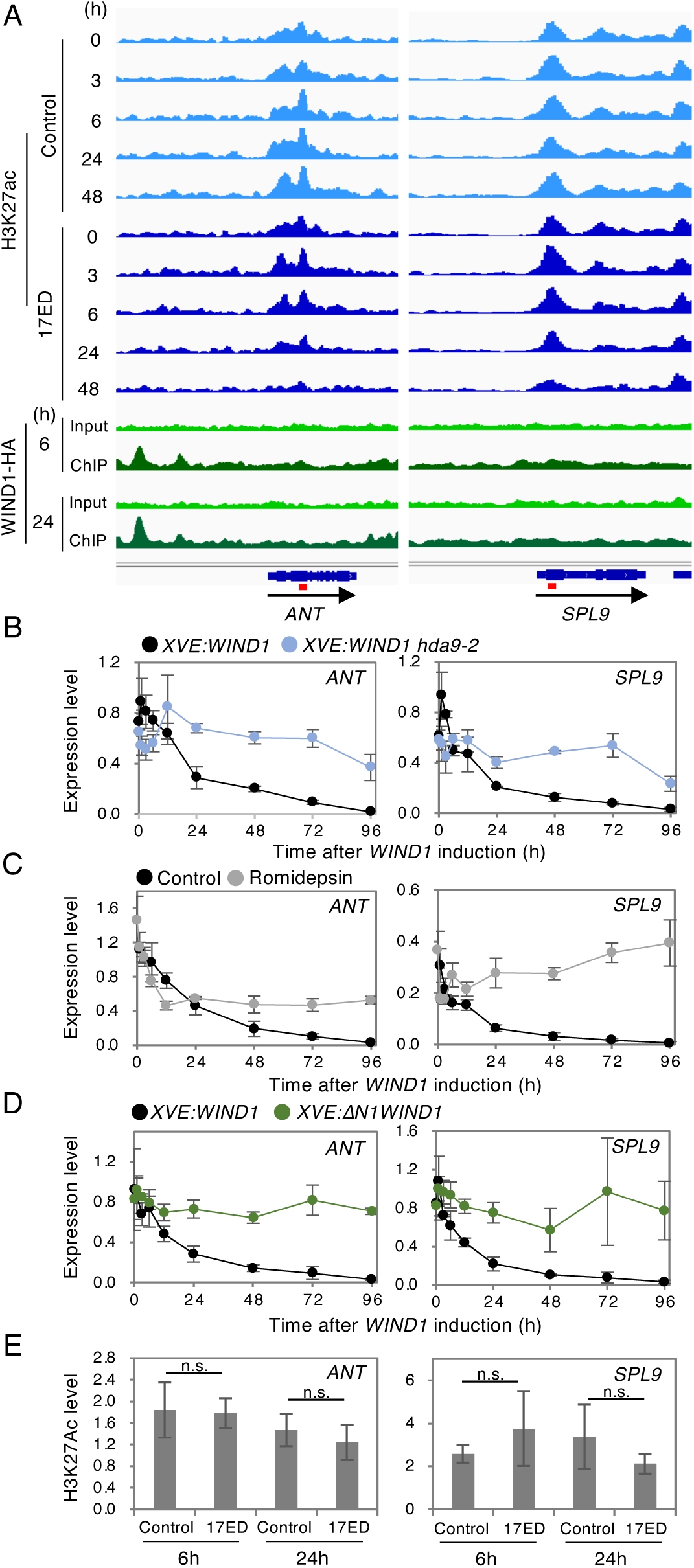
WIND1 downregulates *ANT* and *SPL9* expression in an HDA9-dependent manner. (A) Temporal dynamics of H3K27Ac levels (upper panel) and WIND1 binding (lower panel) at the *ANT* and *SPL9* loci after WIND1 induction. Chromatin immunoprecipitation using an anti-H3K27Ac antibody was performed using *XVE:WIND1* seedlings treated with DMSO (control; light blue) or 17ED (blue) for 0, 3, 6, 24, and 48 h. Chromatin immunoprecipitation using an anti-HA antibody was conducted using *XVE:WIND1-HA* seedlings treated with 17ED for 6 and 48 h. Red lines within the gene structure diagrams indicate the position of primers used for ChIP-qPCR analysis in (E). (B to D) RT-qPCR analysis of *ANT* and *SPL9* expression in *XVE:WIND1* and *XVE:WIND1 hda9-2* (B), in the absence or presence of 5 µM romidepsin (C), and *XVE:WIND1* and *XVE:ΔN1WIND1* at 0, 1, 3, 6, 12, 24, 48, 72, and 96 h after 17ED treatment. Gene expression levels were normalized to *PP2AA3*. Data represent mean ± SE (n = 3 biological replicates). (E) ChIP-qPCR analysis of H3K27Ac levels at the *ANT* and *SPL9* loci in *XVE:ΔN1WIND1* at 6 and 24 hours after 17ED treatment. Statistical significance was assessed by Student’s t-test (n = 3; n.s., not significant).

We further confirmed these interactions using the biomolecular fluorescence complementation (BiFC) analysis with split yellow fluorescent protein (YFP) fusion constructs of WIND1 and HDA9 or ADA2a. As shown in Figures 2D and 2E, co-expression of WIND1 with HDA9 or ADA2a in *N. benthamiana* leaves resulted in YFP fluorescence in the nuclei, indicating that these proteins interact *in planta*. In contrast, fluorescence was strongly reduced when either ΔN1WIND1 or mN1WIND1 was co-expressed, supporting the conclusion that the conserved N1 domain is essential for WIND1’s interaction with HDA9 and ADA2a.

We next examined whether the N1 domain is required for WIND1 function *in planta*. In *XVE:ΔN1WIND1-HA* plants somatic embryo formation was completely abolished upon induction (Figures 2F, 2G, Supplemental Figure 6A). Similarly, while *35S:WIND1-HA* plants exhibited pronounced morphological alterations including callus formation, both *35S:ΔN1WIND1-HA* and *35S:mN1WIND1-HA* plants displayed only mild developmental defects (Supplemental Figures 6C-6E). These results underscore the critical role of the N1 domain in WIND1-mediated cell fate reprogramming.

**Figure 6.**
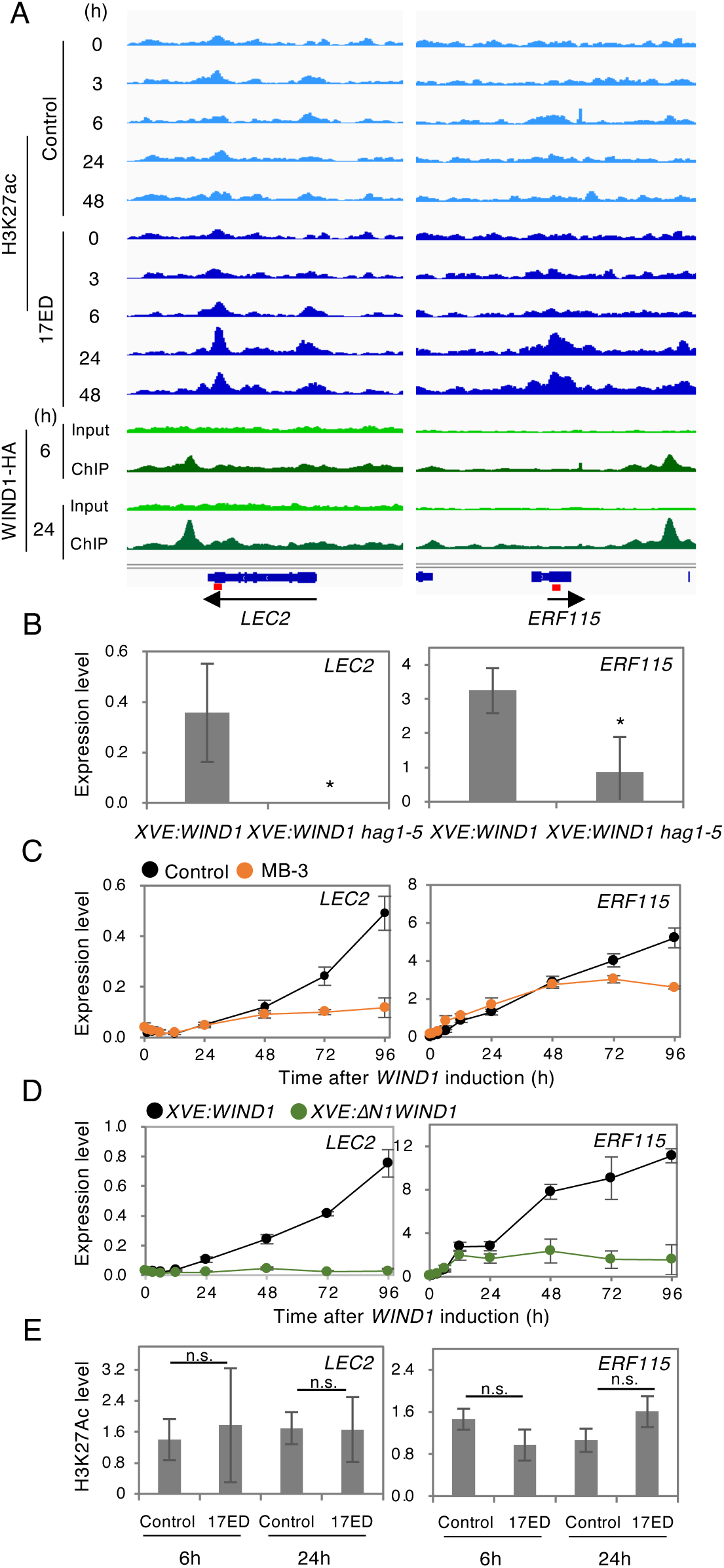
WIND1 upregulates *LEC2* and *ERF115* expression in a HAG1-dependent manner. (A) Temporal dynamics of H3K27Ac levels (upper panel) and WIND1 binding (lower panel) at the *LEC2* and *ERF115* loci after WIND1 induction. Chromatin immunoprecipitation using an anti-H3K27Ac antibody was performed using *XVE:WIND1* seedlings treated with DMSO (control; light blue) or 17ED (blue) for 0, 3, 6, 24, and 48 h. Chromatin immunoprecipitation using an anti-HA antibody was conducted using *XVE:WIND1-HA* seedlings treated with 17ED for 6 and 48 h. Red lines within the gene structure diagrams indicate the position of primers used for ChIP-qPCR analysis in (E). (B) RT-qPCR analysis of *LEC2* and *ERF115* expression in *XVE:WIND1* and *XVE:WIND1 hag1-5* at 48 h after 17ED treatment. Gene expression levels were normalized to *PP2AA3*. Data represent mean ± SE (n = 3 biological replicates). (C-D) RT-qPCR analysis of *LEC2* and *ERF115* expression in *XVE:WIND1* in the absence (control) or presence of 100 µM MB-3 (C) and in *XVE:WIND1* and *XVE:ΔN1WIND1* at 0, 1, 3, 6, 12, 24, 48, 72, and 96 h after 17ED treatment (D). Gene expression levels were normalized to *PP2AA3*. Data represent mean ± SE (n = 3 biological replicates). (E) ChIP-qPCR analysis of H3K27Ac levels at the *LEC2* and *ERF115* loci in *XVE:ΔN1WIND1* at 6 and 24 h after 17ED treatment. Statistical significance was assessed by Student’s t-test (n = 3; n.s., not significant).

### WIND1 selectively modulates H3K27 deacetylation and acetylation across the Arabidopsis genome

Having established a molecular connection between WIND1 and histone acetylation enzymes, we next investigated how WIND1 modulates histone acetylation dynamics. We performed chromatin immunoprecipitation sequencing (ChIP-seq) analysis to profile acetylation of histone H3 at lysine 27th (H3K27Ac) at 0, 3, 6, 24, and 48 h after WIND1 activation in *XVE:WIND1* plants. Across this time course, we identified 3,312 genes exhibiting decreased H3K27 acetylation and 5,210 genes showing increased H3K27Ac levels. These differentially acetylated genes were classified into 11 distinct groups based on the timing and pattern of H3K27Ac changes (Figure 3A). Groups 1-4 included genes with transient or sustained decrease in H3K27Ac while groups 7-10 contained genes with increased acetylation. Group 5 genes initially lost H3K27Ac but regained it at later time points and group 6 displayed the opposite trend. Group 11 genes exhibited non-specific patterns in acetylation over time.

Gene Ontology (GO) analysis revealed that genes extracted from groups 1-4 (Fold change <0.7, *p* <0.05, compared to 0 h) were significantly enriched for biological processes such as shoot system development and photosynthesis (Figure 3B). This is consistent with our earlier finding that HDAC activity is required during the early phase of *WIND1* activation to repress shoot identity (Supplemental Figures 2C and 2E). In contrast, genes from groups 7-10 (Fold chang >1.3, *p* <0.05, compared to 0 h) were enriched for terms related to wound response, defense response, and regulation of embryonic development (Figure 3B, Supplemental Tables 1 and 2). These terms agree with the group of genes upregulated by WIND1 (Iwase et al., 2021), reinforcing the positive correlation between histone acetylation and transcriptional activation.

### WIND1 directly regulates acetylation-coupled gene expression

To further examine the relationship between histone acetylation and WIND1-mediated transcriptional regulation, we conducted RNA-seq analysis on *XVE:WIND1* plants at 0, 1, 3, 6, 12, 24, 48, 72, and 96 h after *WIND1* induction. In total, we identified 6,615 genes that were downregulated and 4,586 genes that were upregulated in response to WIND1 (Supplemental Tables 3 and 4). When these transcriptional profiles were compared with our H3K27ac ChIP-seq data (Supplemental Tables 1 and 2), we observed a statistically significant overlap between genes with decreased H3K27 acetylation and those with reduced expression following *WIND1* activation (Fisher’s exact test; *p* <2.2e-16, odds ratio = 2.6) (Figure 4A). Likewise, genes with increased H3K27Ac levels significantly overlapped with genes showing elevated expression (Fisher’s exact test; *p* <2.2e-16, odds ratio = 2.8) (Figure 4A). These results indicate that WIND1-mediated transcriptional regulation is tightly coupled to changes in histone acetylation status, supporting the idea that WIND1 functions as a chromatin-linked transcriptional regulator.

The 1,258 genes showing decreased H3K27 acetylation and gene expression included key regulators of shoot and floral organ identity such as *AINTEGUMENTA* (*ANT*) and *YABBY5* (*YAB5*), as well as several cell cycle regulators, including *CYCLIN D1;1* (*CYCD1;1*) and *KIP-RELATED PROTEIN 7* (*KRP7*), and photosynthesis-related genes such as *GENOME UNCOUPLED 4* (*GUN4*) and *CHLOROPHYLL A/B BINDING PROTEIN 1* (CAB1). There were also several hormonal regulators like the cytokinin response repressor *SQUAMOSA PROMOTER BINDING PROTEIN-LIKE 9* (*SPL9*) (Figure 4B) (Mizukami and Fischer, 2000; Sarojam et al., 2010; Anzola et al., 2010; Adhikari et al., 2011; Collins et al., 2015; Zhang et al., 2015). Conversely, the 1,641 genes exhibiting both increased H3K27ac and upregulated expression included multiple regulators of somatic embryogenesis, callus formation and shoot regeneration such as *LEC2, ETHYLENE RESPONSE FACTOR 115* (*ERF115*)*, RELATED TO AP2 6-LIKE* (*RAP2.6L*) and *WUSCHEL-RELATED HOMEOBOX 13* (*WOX13*) (Stone et al., 2001; Che et al., 2006; Heyman et al., 2016; Iwase et al., 2021; Ikeuchi et al., 2022), as well as genes involved in tracheary element formation and defense responses such as *VASCULAR-RELATED NAC-DOMAIN PROTEIN 7* (*VND7*), and *RAP2.6* (Figure 4B) (Kubo et al., 2005; Ali et al., 2013). These results suggest that WIND1 plays a dual role in cell fate reprogramming by actively repressing pre-existing developmental programmes while promoting the acquisition of new cellular identities.

To determine whether WIND1 directly regulates the expression of its downstream genes, we generated *XVE:WIND1-HA* plants and performed ChIP-seq analysis using anti-HA antibody to identify genome-wide WIND1 biding sites. To capture early binding events, potentially preceding or coinciding with transcriptional changes, we selected 2 time points, 6 and 24 h after WIND1 induction. In total, we identified 3,235 WIND1 binding sites across these time points (Supplemental Tables 5 and 6). Motif analysis of these regions revealed that WIND preferentially binds to GC-rich G(T/A)CGG(T/C) motifs and their complementary sequences (G/A)CCG(T/A) C at both 6h and 24h post induction (Supplemental Figure 7A). These binding preferences are consistent with an earlier report that the AP2/ERF DNA binding domain recognizes the GCC box motif ((A/G)CCGNC) (Ohme-Takagi and Shinshi, 1995), confirming that our ChIP-seq captured genuine WIND1-DNA interactions. Among the 1,641 genes that showed both increased H3K27Ac levels and elevated expression following WIND1 induction, 616 genes (37.5%) were directly bound by WIND1 (Figure 4A). In contrast, of the 1,258 downregulated genes with decreased H3K27Ac, only 114 genes (9.1%) exhibited direct WIND1 binding (Figure 4A). Consistent with this trend, we observed direct WIND1 binding at the loci of several key WIND-induced reprogramming genes but only at a subset of WIND1-repressed genes (Figure 4B). These results suggest that WIND1 primarily functions as a direct activator of gene expression, while its repressive effects are more often indirect or mediated through additional regulatory factors. To further explore the basis of this differential regulatory behaviour, we performed separate motif enrichment analyses on the 616 directly activated genes and the 114 directly repressed genes. This revealed a clear difference in the enriched DNA motifs between the two groups (Supplemental Figure 7B). While the canonical GCC-box motif was most strongly enriched among the directly activated targets, the repressed loci were instead enriched for a distinct sequence motif, (A/C)CACA. These findings suggest that WIND1 may engage distinct DNA sequence contexts when functioning as a transcriptional repressor, potentially reflecting different modes of chromatin engagement or cofactor recruitment.

### WIND1 suppresses *ANT* and *SPL9* expression in an HDA9-dependent manner

Among the genes repressed by WIND1, the ANT transcription factor plays a crucial role in leaf development (Mizukami and Fischer, 2000), while SPL9 negatively regulates cytokinin signalling by repressing the expression of type-B Arabidopsis response regulator (ARR) genes (Zhang et al., 2015). We therefore selected these two genes to further explore the mechanisms by which WIND1 represses target gene expression. As shown in Figures 4B, 5A, 5B and Supplemental Table 1, both *ANT* and *SPL9* loci displayed a progressive decline in H3K27 acetylation and transcript abundance following 6 h of WIND1 induction. Notably, ChIP-seq analysis detected direct WIND1-HA binding at the *ANT* locus but not at *SPL9* at either 6 or 24 h post induction (Figure 5A), implying that these genes are repressed *via* at least partially distinct regulatory mechanisms.

To determine whether HDA9 is required for WIND1-mediated repression of *ANT* and *SPL9*, we compared their expression levels in *XVE:WIND1* and *XVE:WIND1 hda9-2* plants by RT-qPCR. As shown in Figure 5B, expression of both genes remained relatively high in the *XVE:WIND1 hda9-2* background compared to *XVE:WIND1* alone. Similarly, in *XVE:WIND1* plants treated with romidepsin, expression of *ANT* and *SPL9* was sustained for up to 96 h following WIND1 induction (Figure 5C), further supporting a role for HDA9 in their repression.

We next investigated whether the N1 domain of WIND1 is required for this transcriptional repression by comparing gene expression between *XVE:WIND1* and *XVE:ΔN1WIND1* plants. As expected, expression of *ANT* and *SPL9* remained elevated in *XVE:ΔN1WIND1* (Figure 5D), clearly demonstrating the essential role of the N1 domain in this process. Importantly, ChIP-qPCR analysis using an anti-H3K27ac antibody revealed that H3K27 acetylation levels at these loci remain unchanged in *XVE:ΔN1-WIND1* plants (Figure 5E). Together, these findings suggest that WIND1 represses *ANT* and *SPL9* expression through its interaction with HDA9 *via* the N1 domain, likely by promoting histone deacetylation at their genomic loci.

### WIND1 activates *LEC2* and *ERF115* expression in a HAG1-dependent manner

To investigate the mechanism of gene activation by WIND1, we focused on *LEC2* and *ERF115*, two key genes upregulated by WIND1. *LEC2* promotes somatic embryogenesis when overexpressed (Stone et al., 2001), while *ERF115* functions as a central regulator of wound-responsive cell reprogramming (Heyman et al., 2016). ChIP-seq analysis revealed that WIND1-HA binds directly at these loci as early as 6 h after induction and H3K27 acetylation at these loci continue progressively increases up to 24 h post induction (Figures 4B, 6A, Supplemental Table 2).

Consistent with these findings, RT-qPCR analysis confirmed that *LEC2* and *ERF115* transcript levels were upregulated following *WIND1* induction (Figure 6B). Importantly, this upregulation was significantly impaired in the *XVE:WIND1 hag1-5* background and *XVE:WIND1* plants treated with MB-3 (Figures 6B and 6C), indicating that HAG1-dependent histone acetylation is required for WIND1-mediated activation of these genes. Furthermore, in *XVE:ΔN1WIND1* plants treated with 17ED, *LEC2* and *ERF115* expression remained low, and this was accompanied by unchanged H3K27 acetylation levels at these loci (Figures 6D and 6E). These results together support that WIND1 activates *LEC2* and *ERF115* expression by HAG1-mediated histone acetylation, which is dependent on the N1 domain of WIND1.

To test whether LEC2 and ERF115 are involved in WIND-induced somatic embryogenesis, we introduced the *lec2* and *erf115* mutations into *XVE:WIND1* plants by genetic crossing. As shown in Figures 7A-D, somatic embryo formation was completely abolished in *XVE:WIND1 lec2* plants whereas *XVE:WIND1 erf115* plants still formed embryos. These data demonstrate that LEC2 is essential for WIND1-mediated somatic embryogenesis while ERF115 is dispensable in this biological context.

**Figure 7.**
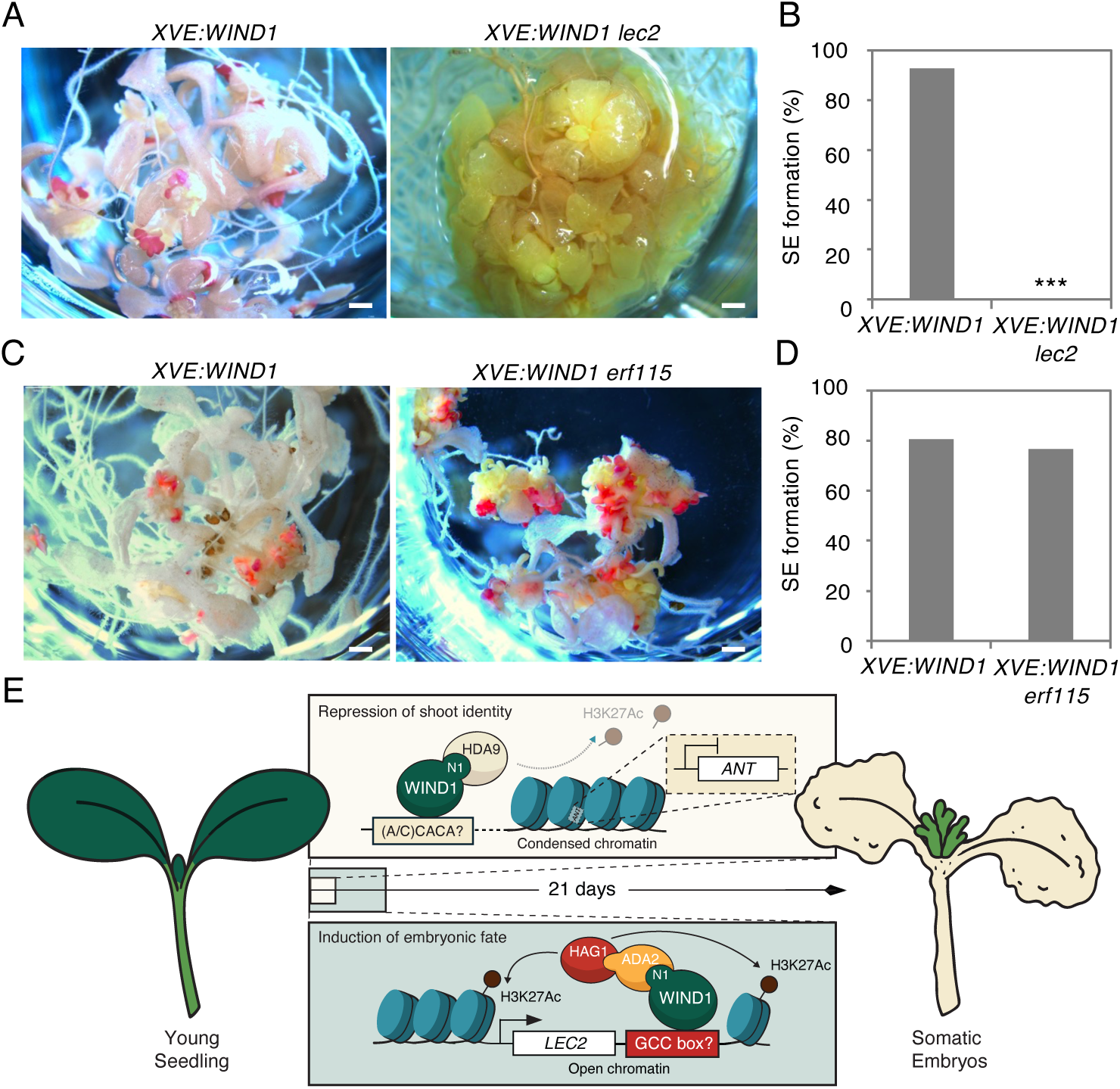
LEC2 is required for WIND1-induced somatic embryogenesis. (A and B) Sudan Red-stained somatic embryos on 17ED-treated *XVE:WIND1* and *XVE:WIND1 lec2* seedlings (A). Quantitative data of SE formation (B). Scale bars, 1 mm. Statistical significance was determined by proportion test (n = 69 for *XVE:WIND1* and 59 for *XVE:WIND1 lec2*. \*\*\**p* < 2.2e-16). (C and D) Sudan Red-stained somatic embryos on 17ED-treated *XVE:WIND1* and *XVE:WIND1 erf115* seedlings (C). Quantitative data of SE formation (D). Scale bars, 1mm. Statistical significance was determined by proportion test (n = 31 for *XVE:WIND1* and 30 for *XVE:WIND1 erf115*). (E) Proposed model for the dual function of WIND1 in somatic embryogenesis. WIND1 promotes HDA9-mediated histone deacetylation at target loci, such as *ANT*, to repress the expression of genes associated with pre-existing shoot identity. In parallel, WIND1 induces HAG1-dependent histone acetylation at loci such as *LEC2* to activate genes required for acquiring embryonic identity. This coordinated loss of original cell fate and activation of embryonic programs are essential for successful somatic embryogenesis.

## DISCUSSION

In this study we demonstrate that WIND1 promotes somatic embryogenesis through a dual chromatin-based mechanism involving histone deacetylation and acetylation (Figure 7E). Our data show that WIND1 physically interacts with the HDA9 and ADA2a-HAG1 module using its conserved N-terminal domain (Figures 2 and Supplemental Figures 3-5). These interactions allow WIND1 to repress genes associated with shoot identity and function through HDA9-dependent histone deacetylation while simultaneously activating key reprogramming regulators *via* HAG1-mediated histone acetylation (Figures 3 and 4). Genetic and pharmacological perturbations further confirm that both repression of the pre-existing developmental programme and activation of embryonic regulators are essential for successful cell fate reprogramming (Figures 1, 6, 7 and Supplemental Figure 2). Although the interplay between transcription factors and chromatin modifiers is well established, the capacity of a single transcription factor to recruit both repressive and activating chromatin regulators remains underexplored in plants and has been only rarely described in animals. One notable example is the muscle differentiation factor (MyoD), which interacts with HDACs to repress fibroblast identity genes while also recruiting HATs to activate muscle-specific genes, thereby coordinating cell fate transitions (Puri and Sartorelli, 2000; Mal and Harter, 2003). Our findings suggest that WIND1 similarly acts as a bifunctional chromatin regulator, coupling the repression of existing identity with activation of new developmental programmes. Intriguingly, a recent study in yeast revealed that many transcription factors possess both activating and repressing capacities, although the underlying molecular mechanisms are not yet understood (Mahendrawada et al., 2025). Our results raise the possibility that the ability to recruit both HDACs and HATs may represent a mechanism that underlies such dual functionality in transcriptional regulation.

An important unsolved question is how the WIND1-HDA9 and WIND1-ADA2-HAG1 modules selectively target different genomic loci to impose opposing chromatin modifications. One possibility is that WIND1 recognizes distinct regulatory *cis* elements at deacetylated versus acetylated loci, thereby recruiting specific chromatin-modifying complexes in a sequence-specific manner. Alternatively, these modules may interact with additional transcriptional cofactors that confer locus-specific targeting. Such combinatorial interactions could help determine whether a WIND1-bound locus is destined for transcriptional activation or repression. Consistent with these ideas, the motif enrichment analyses of our ChIP-seq datasets uncovered sequence elements that are uniquely associated with repressed or activated targets (Supplemental Figure 7). Further characterisation of these specificity determinants will be critical for understanding how WIND1 integrates chromatin context with transcriptional control during cell fate transitions.

It is also notable that our ChIP-seq analyses detected WIND1 binding at only 9.1% of genes that were deacetylated and transcriptionally repressed following WIND1 induction (Figure 4), implying that many of these genes may be regulated indirectly. WIND1, for example, may activate secondary transcriptional repressors that in turn downregulate these targets. Alternatively, WIND1 might bind to distal regulatory elements that were not captured under our experimental conditions. Recent studies, including Mahendrawada et al. (2025) have predicted the importance of long-range chromatin interactions in defining transcriptional outcomesc (Mahendrawada et al., 2025), which may not be evident in conventional peak-calling approaches. Another possibility is that WIND1 binding at repressed loci is transient or occurring at earlier time points than those sampled in this study. These scenarios highlight the importance of high-temporal resolution assays and 3D chromatin conformation mapping to fully characterize the dynamics of WIND1-mediated transcriptional repression.

Our data also reveal a subset of WIND1 responsive genes undergo transcriptional changes independent of detectable changes in H3K27 acetylation (Figure 4). These include both up- and down-regulated genes, suggesting that WIND1 can also regulate transcription through acetylation-independent mechanisms. The existence of these acetylation-independent modes highlights the multifaceted nature of WIND1-mediated transcriptional reprogramming and suggests that WIND1 acts as a central integrator of diverse regulatory inputs during cell fate transitions. Future studies dissecting these additional layers of regulation will further deepen our understanding of how transcription factors coordinate complex developmental reprogramming events.

## METHODS

### Plant materials and growth condition

All Arabidopsis plants used in this study were in the Col-0 background. *XVE:WIND1*, *WIND1pro:WIND1-SRDX*, *hag1-1* (SALK_150784), *hag1-5* (SALK_048427), *hag3-1* (GABI_555H06), *hag3-2* (SALK_060819)*, hda6-6* (*axe1-5*), *hda9-1* (SALK_007123)*, hda9-2* (GABI_305G03), *hda19-3* (SALK_139445)*, hda19-5* (CRISPR/CAS9), *lec2* (SALK_015228), *erf115* (SALK_021981) were described previously (Iwase et al., 2011; Heyman et al., 2013; Ikeuchi et al., 2015; Rymen et al., 2019; Iwase et al., 2021; Hung et al., 2023; Temman et al., 2023). T-DNA insertion lines were obtained from the Arabidopsis Biological Resource Center (ABRC). Plants were grown on 0.6% (w/v) gelzan plates containing Murashige and Skoog (MS) salt and 1% sucrose medium at 22°C with a photoperiod of 16 h white light and 8 h darkness, unless noted otherwise. For *WIND1* induction, *XVE:WIND1* plants were germinated and grown in 1/2 MS liquid medium with rotation (100rpm). 5-day-old seedlings were treated with DMSO or 1 µM 17-β-estradiol (Sigma-Aldrich). γ-butyrolactone (MB-3, CAS No. 778649-18-6, Sigma), romidepsin (FK228, CAS No. 128517-07-7, Sigma), and Ky-2 (Nishino et al., 2004) were applied to the 1/2 MS liquid medium.

### Plasmid construction and transgenic plants

To construct the *p35S:WIND1-HA and p35S:ΔN1WIND1-HA* vectors, the protein coding region of *WIND1* with HA tag was PCR amplified from the *pBCKH-35S:WIND1* vector (Iwase et al., 2011) and cloned into the *p35SG* vector (Ohta et al., 2001). Resulting fragments were subcloned into the *pBCKK* vector for plant transformation (Mitsuda et al., 2005). The *p35S:mN1WIND1-HA* vector was constructed by site-directed mutagenesis using the *p35S:WIND1-HA* vector as a template and primers designed to introduce the mutation. To construct the *XVE:ΔN1WIND1-HA* vector, the *ΔN1WIND1* fragment was PCR amplified from the *p35S:ΔN1WIND1-HA* vector and cloned into the *pER8* vector (Zuo et al., 2000). For plant transformation, T-DNA vectors carrying an appropriate construct were introduced into *Agrobacterium tumefaciens* strain GV3101 by electroporation and Arabidopsis plants were transformed by the floral dip method (Clough and Bent, 1998). The primer sequences used for generating these constructs are listed in Supplemental Table 7.

### Yeast two-hybrid (Y2H) assay

The yeast two-hybrid experiments were performed with the Matchmaker Gold Yeast Two-Hybrid system (Takara Bio) according to the manufacturer’s instructions. To test the interaction between WIND1 and candidate proteins, coding region of *WIND1*, *HDA6, HDA9*, *HDA19*, *ADA2a*, *ADA2b*, and *HAG1* were amplified from cDNA by PCR, cloned into the *pDONR207* entry vector and then subcloned into the *pDEST-GAKT7* and *pDEST-GBKT7* vectors by the Gateway LR reaction (Thermo Fisher Scientific). The yeast strain Y2HGold was co-transformed with these vectors, grown on synthetic dextrose (SD)/- Leu-Trp media. Transactivation activity was further evaluated based on the colony formation on SD/-Trp-Leu-Ade-His media containing 0.2 mg/L of Aureobasidin A (+AbA). Since WIND1 C-terminal region has an autoactivation activity in the yeast cells, full-length or truncated WIND1 bait construction lacking C-terminal region from 318^th^ to the last 334^th^ amino acids (ΔC2WIND1) was used. The primer sequences used for cloning are listed in Supplemental Table 7.

### Co-immunoprecipitation (Co-IP) assay

Full-length cDNA of WIND1, ΔN1WIND1, and mN1WIND1 were fused with the HA tag by PCR and then cloned into the *pBCKK* vector. ADA2a and HDA9 cDNAs were fused with the FLAG tag by PCR and then introduced into the *pEAQ-HT-DEST1* vector (Sainsbury et al., 2009). The resulting *WIND1-HA* plasmids were co-injected with *ADA2a-FLAG* or *HDA9-FLAG* plasmids into *N. benthamiana* leaves by agroinfiltration. The following Co-IP assay was performed as described previously (Shibata et al., 2022). The primer sequences used for the cloning are listed in Supplemental Table 7.

### Biomolecular fluorescence complementation (BiFC) assay

For the BiFC assays, full-length, truncated and mutated cDNAs of WIND1 (Full, ΔN1, and mN1), ADA2a, and HDA9 were PCR amplified, cloned into the *pDONR207 or pDONR221* vector (Invitrogen) and recombined into the *pEarleyGate201-YN* and *pEarleyGate202-YC* vectors (Hung et al., 2023). These vectors were transiently transformed into *N. benthamiana* leaves by *A. tumefaciens* strain AGL-1. Signals of fluorescence proteins were visualized by a Leica TCP SP5 II confocal laser microscope. The primer sequences used for cloning are listed in Supplemental Table 7.

### Somatic embryo formation assay

5-day-old *XVE:WIND1* seedlings were treated with DMSO or 1 µM 17-β-estradiol (Sigma-Aldrich) for up to 21 days. Somatic embryos were stained by a protocol previously reported (Ikeuchi et al., 2015) and observed under Leica M165 C stereomicroscope. The hormone-induced somatic embryogenesis assay from immature embryos was performed as described previously (Gaj, 2001; Wang et al., 2020).

### Callus formation assay

The callus formation assay from petioles was performed as described previously (Iwase et al., 2017). Callus were observed under Leica M165 C stereomicroscope.

### RNA isolation, quantitative RT-qPCR and RNA-seq

RT-qPCR was performed as previously described (Iwase et al., 2021). Total RNA was isolated from 17-β-estradiol-treated whole seedlings of *XVE:WIND1* plants for each chemical treatment experiment (DMSO, MB-3 and romidepsin) and time point (0, 1, 3, 6, 12, 24, 48, 72 and 96 h) with RNeasy Plant Mini Kit (QIAGEN) according to manufacturer’s instruction. Extracted RNA was reverse transcribed by a PrimeScript RT-PCR kit with DNase I (Takara) in accordance with the accompanying protocol. Transcript levels were determined by RT-qPCR using a THUNDERBIRD SYBR qPCR Mix kit (Toyobo) and an Mx399P QPCR system (Agilent). For each sample, the expression levels were quantified for 3 biological replicates and normalised to those of the *PROTEIN PHOSPHATASE 2A SUBUNIT A3* (*PP2AA3*) gene. For RNA sequencing, isolated RNA was subjected to library preparation using a KAPA stranded mRNA sequencing kit (Roche) with NEBNext Multiplex Oligos for Illumina (New England Biolabs) as adapters and Agencourt AMPure XP (Beckman Coulter) beads in place of KAPA Pure Beads. Single-end sequencing was performed on an Illumina NextSeq500 platform. Mapping was carried out using Bowtie version 0.12.9 (Langmead and Salzberg, 2012). On average, 77% (43 to 85%) of reads were mapped to the Arabidopsis TAIR10 Arabidopsis genome. DEGs were identified using the edgeR package(Robinson et al., 2009) in R/Bioconductor (https://www.r-project.org/) after normalization of total read counts with the trimmed mean of M-values method. DEGs were defined as those that showed |log2FC| > 2 in transcript levels (p < 0.01 and FDR < 0.05). All primer sequences are listed in Supplemental Table 7.

### Enhanced chromatin immunoprecipitation (eChIP)-seq

The eChIP method was applied for both H3K27ac and WIND1-HA binding detection according to Mori et al. with several modifications (Mori et al., 2023). Approximately 100 mg (for H3K27ac) or 500 mg (for WIND1-HA) of aerial tissues from seedlings grown in liquid culture were ground in liquid nitrogen and added to the fixation buffer, i.e. phosphate buffered saline (PBS) with 1% formaldehyde, 1 mM Pefabloc SC (Roche) and cOmplete protease inhibitor cocktail (Roche), and incubated at room temperature for 10 min. Glycine was added to a concentration of 0.2 M and further incubated at room temperature for 5 min, after which the supernatant was removed by centrifugation. Pellets were lysed in 180 μl of Buffer S (50 mM HEPES-KOH (pH 7.5), 150 mM NaCl, 1 mM EDTA, 1% Triton X-100, 0.1% sodium deoxycholate, 1% SDS) for 10 min at 4°C. The homogenate was mixed with 720 μl of Buffer F (50 mM HEPES-KOH (pH 7.5), 150 mM NaCl, 1 mM EDTA, 1% Triton X-100, 0.1% sodium deoxycholate). The chromatin was fragmented into 200∼600 bp by sonication using Covaris (Covaris). The antibodies used were ab1791 (Abcam) for H3, ab4729 (Abcam) for H3K27Ac and ab9110 (Abcam) for HA tag. For ChIP sequencing, isolated DNA was subjected to library preparation using a KAPA Hyper Prep Kit (Roche) with NEBNext Multiplex Oligos for Illumina (New England Biolabs) as adapters and Agencourt AMPure XP (Beckman Coulter) beads in place of KAPA Pure Beads. Pair-end sequencing was performed on an Illumina Novaseq 6000 platform. The raw sequence data were processed using Bowtie2 to map the reads to the Arabidopsis genome (TAIR10) (Langmead and Salzberg, 2012). The mapped reads were retained for further analysis. The alignments were first converted to Wiggle (WIG) files using deepTools (Ramírez et al., 2014). The data were then imported into the Integrated Genome Viewer (IGV) for visualization (Robinson et al., 2011). To identify the relative H3K27Ac level, the raw reads of H3 and H3K27Ac within gene bodies were calculated by the deepTools (Ramírez et al., 2014). The reads of H3K27Ac were first normalized to H3 and then normalized to 0-hour untreated sample. GO analyses were performed using PANTHER GO Enrichment Analysis (Mi et al., 2021) (http://geneontology.org) with the PANTHER Overrepresentation Test (Released 20240807). Genes were annotated using the GO database (Released 2025-03-16) and categorised by Biological Processes. The WIND1 binding peaks were analyzed by MACS2 (Feng et al., 2012), and the nearby targeted genes of the peak summits were identified by the ChIPseeker (Yu et al., 2015). To identify DNA motifs enriched sites, 400-bp sequences encompassing each peak summit (200 bp upstream and 200 bp downstream) were extracted and searched for enriched DNA motifs using MEME-ChIP with the default parameters (Machanick and Bailey, 2011).

## Supporting information

Iwase_Supplemental_Tables_to_submit_BioRxiv

## DATA AND CODE AVAILABILITY

Data for ChIP-seq and RNA-seq experiments are available from the National Center for Biotechnology Information BioProject Database under the code GSE301977 for ChIP-seq and GSE302400 for RNA-seq.

## ACKNOWLEDGEMENTS

The authors thank Chika Ikeda, Mariko Mouri, Noriko Doi, Akiko Hanada and Ayami Furuta for their technical assistance. We thank Akihiro Ito and Minoru Yoshida for providing Ky-2 and Nobutaka Mitsuda for the Y2H vectors harbouring PIF3 or HYH.

## AUTHOR CONTRIBUTIONS

A.I. and K.Su conceived and designed the experiments. A.I., A.T, F-Y.H., A.K. Y.C.I performed the experiments and analysed the data. Y.K. and K.Sh helped with Co-IP and S.I. helped with ChIP experiments. T.S. sequenced the RNAseq libraries. A.I. and K.Su wrote the manuscript with contributions from all other authors. All authors revised and approved the final manuscript.

## SUPPLEMENTAL INFORMATION

**Supplemental Figure 1.**
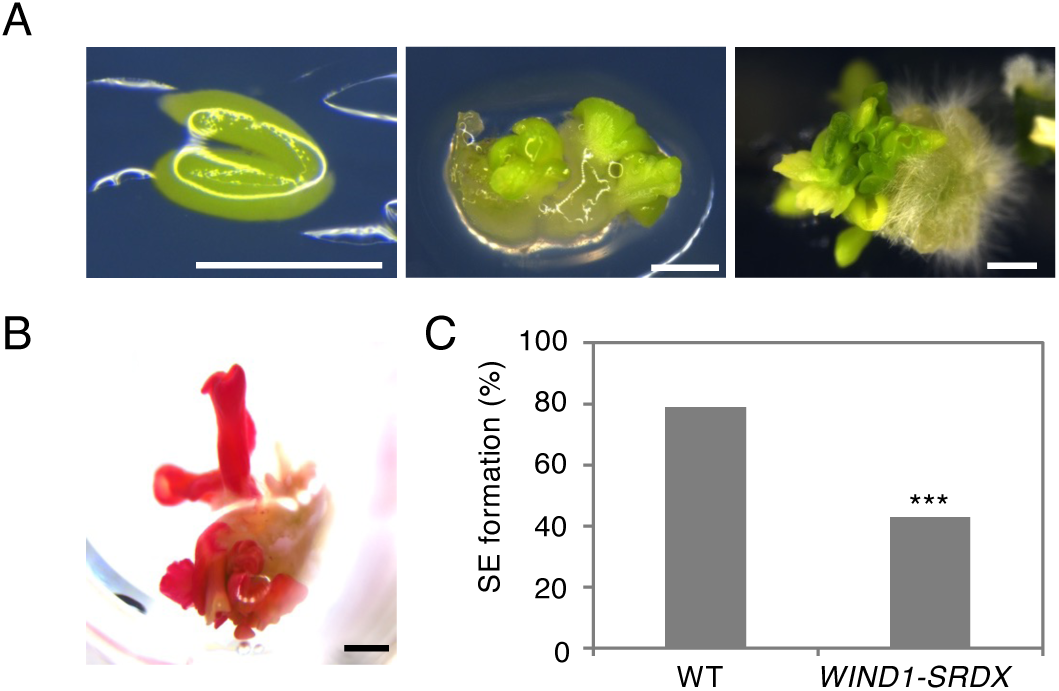
WIND1 is required for the hormone-induced somatic embryogenesis from immature embryos. (A) WT immature embryo (left), embryo-like structures emerging from an immature embryo cultured on 2,4-Dichlorophenoxyacetic acid (2,4-D) containing somatic embryo induction medium for 14 d (middle), and SE-like structures formed on callus cultured on hormone-free medium for 10 d (right). Scale bars, 1mm. (B) Sudan Red-stained somatic embryos. Scale bar, 1mm. (C) Quantitative data of somatic embryo (SE) formation in wild type (WT) and *WIND1-SRDX* plants shown as ratio (%) of seedlings that formed SE among all tested seedlings. Statistical significance was determined by proportion test (n = 137 for WT, 133 for *WIND1-SRDX*, \*\*\**p* < 2.89e-9).

**Supplemental Figure 2.**
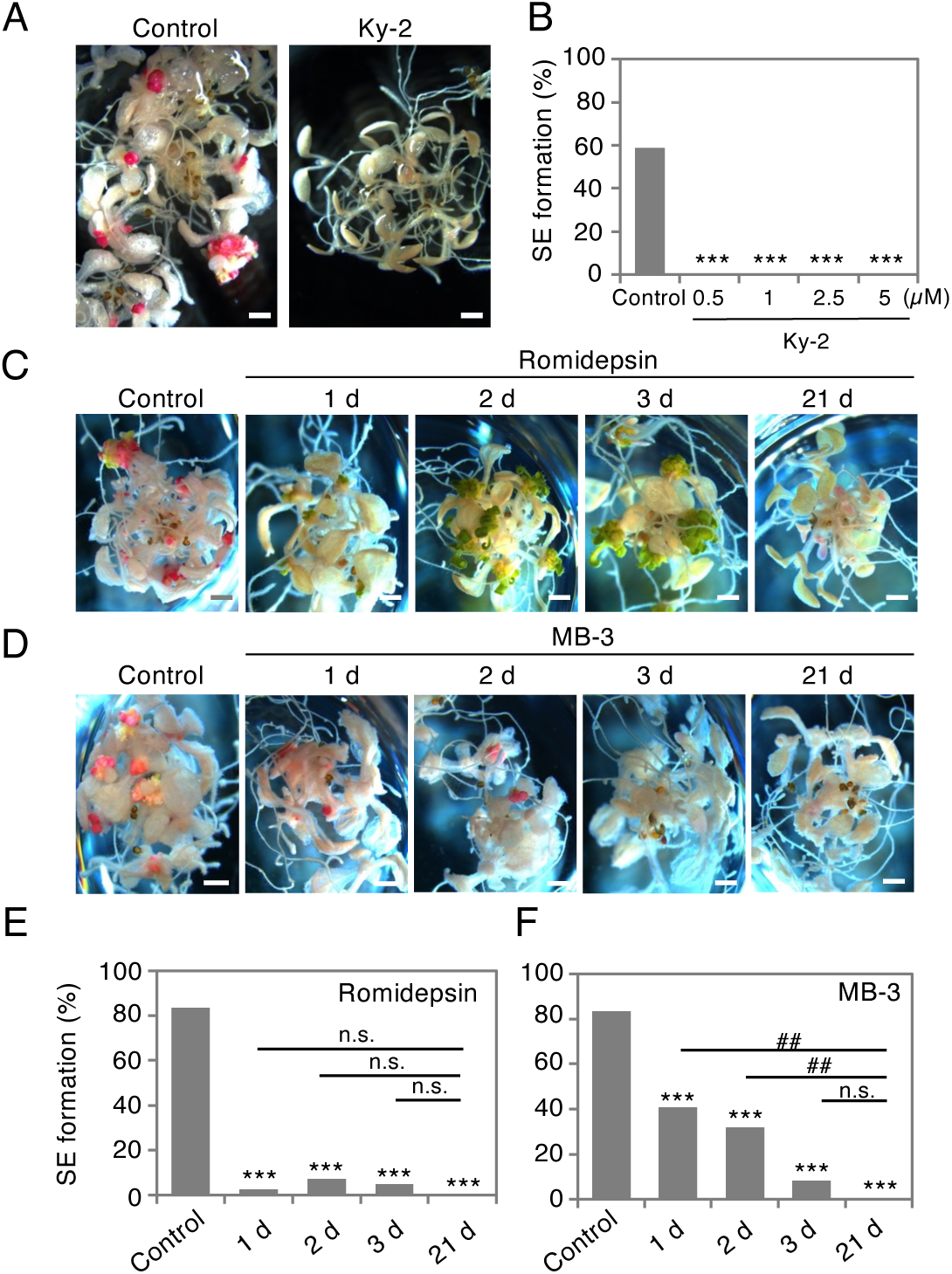
Histone deacetylation and acetylation are both required for WIND1-induced somatic embryogenesis. (A and B) Sudan Red-stained somatic embryos on 17ED-treated *XVE:WIND1* seedlings in the presence of 5 µM Ky-2, a class I histone deacetylase inhibitor (A). Quantitative data of SE formation (B). Scale bars, 1mm. Statistical significance was determined by proportion test (n = 21∼39 for each concentration, \*\*\**p* <1.23e-5). (C) Sudan Red-stained somatic embryos on 17ED-treated *XVE:WIND1* seedlings in the absence (control) or presence of 5 µM romidepsin for 1, 2, 3, or 21 d. Scale bars, 1mm. (D) Sudan Red-stained somatic embryos on 17ED-treated *XVE:WIND1* seedlings in the absence (control) or presence of 100 µM MB-3 for 1, 2, 3, or 21 d. Scale bars, 1mm. (E) Quantitative data of SE formation on 17ED-treated *XVE:WIND1* seedlings in the absence (control) or presence of 5 µM romidepsin for 1, 2, 3, or 21 d. Statistical significance was determined by proportion test (n = 54 for control, 37 for 1 d, 42 for 2 d and 41 for 3 d, and 43 for 21 d, \*\*\**p* < 1.73e-13 compared to control; n.s., no significance compared to 21 d). (F) Quantitative data of SE formation on 17ED-treated *XVE:WIND1* seedlings in the absence (control) or presence of 100 µM MB-3 for 1, 2, 4, or 21 d. Statistical significance was determined by proportion test (n = 54 for control, 32 for 1 d, 41 for 2 d and 37 for 3 d, and 44 for 21 d, \*\*\**p* < 1.19e-4 compared to control; ##*p* < 0.0005 compared to 21 d; n.s., no significance compared to 21 d).

**Supplemental Figure 3.**
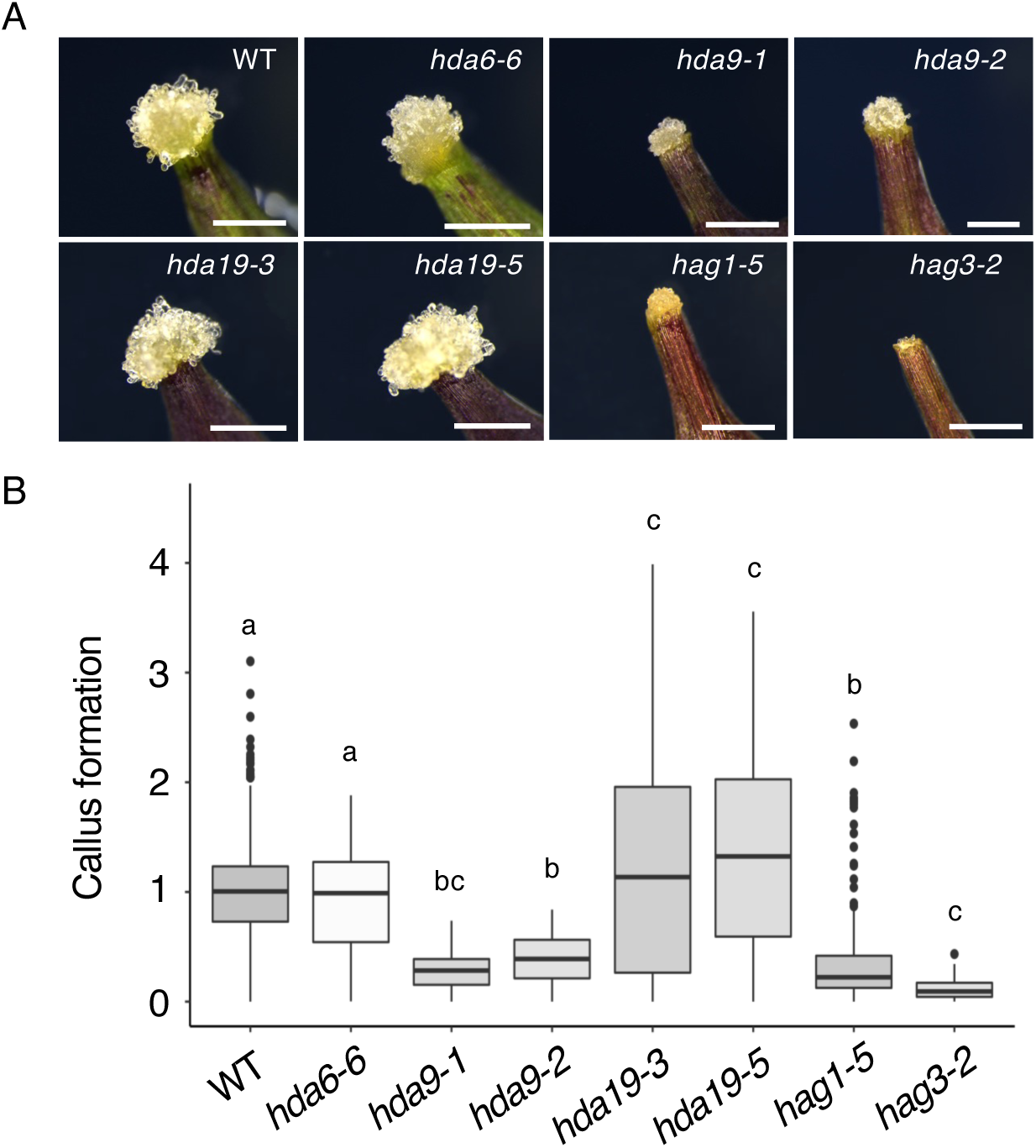
HDA9 and HAG1 are required for wound-induced callus formation. (A) Light micrograph of callus forming at wound sites in WT, *hda6-6*, *hda9-1*, *hda9-2*, *hda19-3*, *hda19-5*, *hag1-5*, and *hag3-2* leaf petioles. Bars, 1mm. (B) Box plots representing the distribution of projected callus area. Horizontal line shows median, the lower and upper bounds of each box plot denote the first and third quartiles, and whiskers above and below the box plot indicate 1.5 times the interquartile range. Outliers are shown as dots. Letters indicate statistical significance determined by ANOVA and Tukey’s multi-comparison test (n = 69 to 539 per genotype, *p* < 0.05).

**Supplemental Figure 4.**
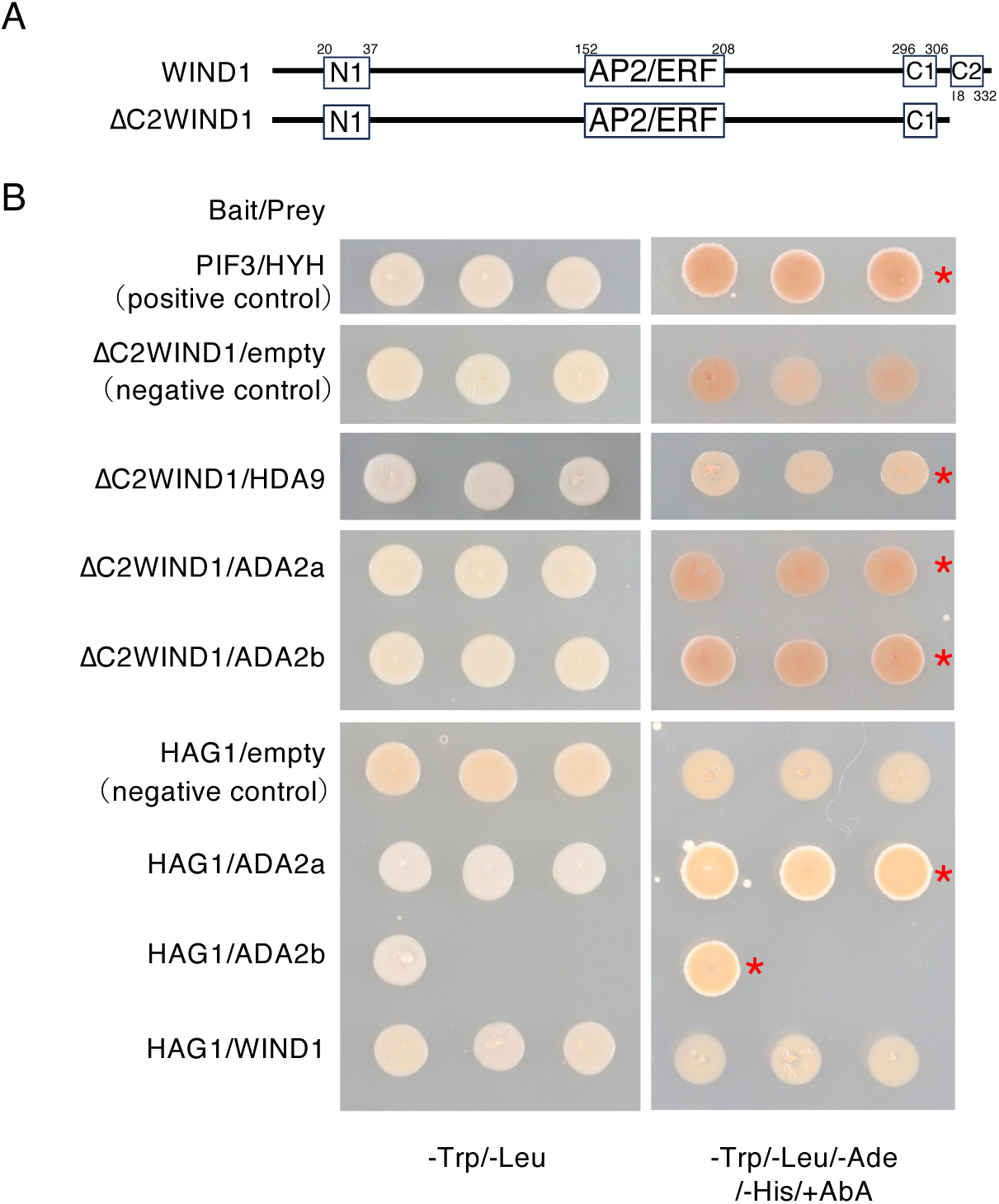
WIND1 interacts with HDA9 and ADA2 proteins in yeast cells. (A) Schematic diagram of *WIND1* constructs. White boxes represent amino acid sequences conserved among WIND1 homologues (Iwase et al., 2013) and numbers above boxes show the order of amino acids. N1; conserved N-terminal domain 1, AP2/ERF; DNA binding domain, C1 and C2; conserved C-terminal domain 1 and 2. WIND1; WIND1 full-length, ΔC2WIND1; WIND1 lacking the C2 domain. (B) Y2H data showing the interactions between WIND1 and HDA6, ADA2a and ADA2b. Transformed yeast cells grown on SD-Trp/-Leu (left) and SD-Trp/-Leu/-Ade /-His/+AbA medium (right). PIF3-HYH binding was used as a positive control (Favero et al., 2020). ΔC2WIND1-empty prey vector and HAG1-empty prey vector served as negative controls. Asterisks indicate positive yeast growth.

**Supplemental Figure 5.**
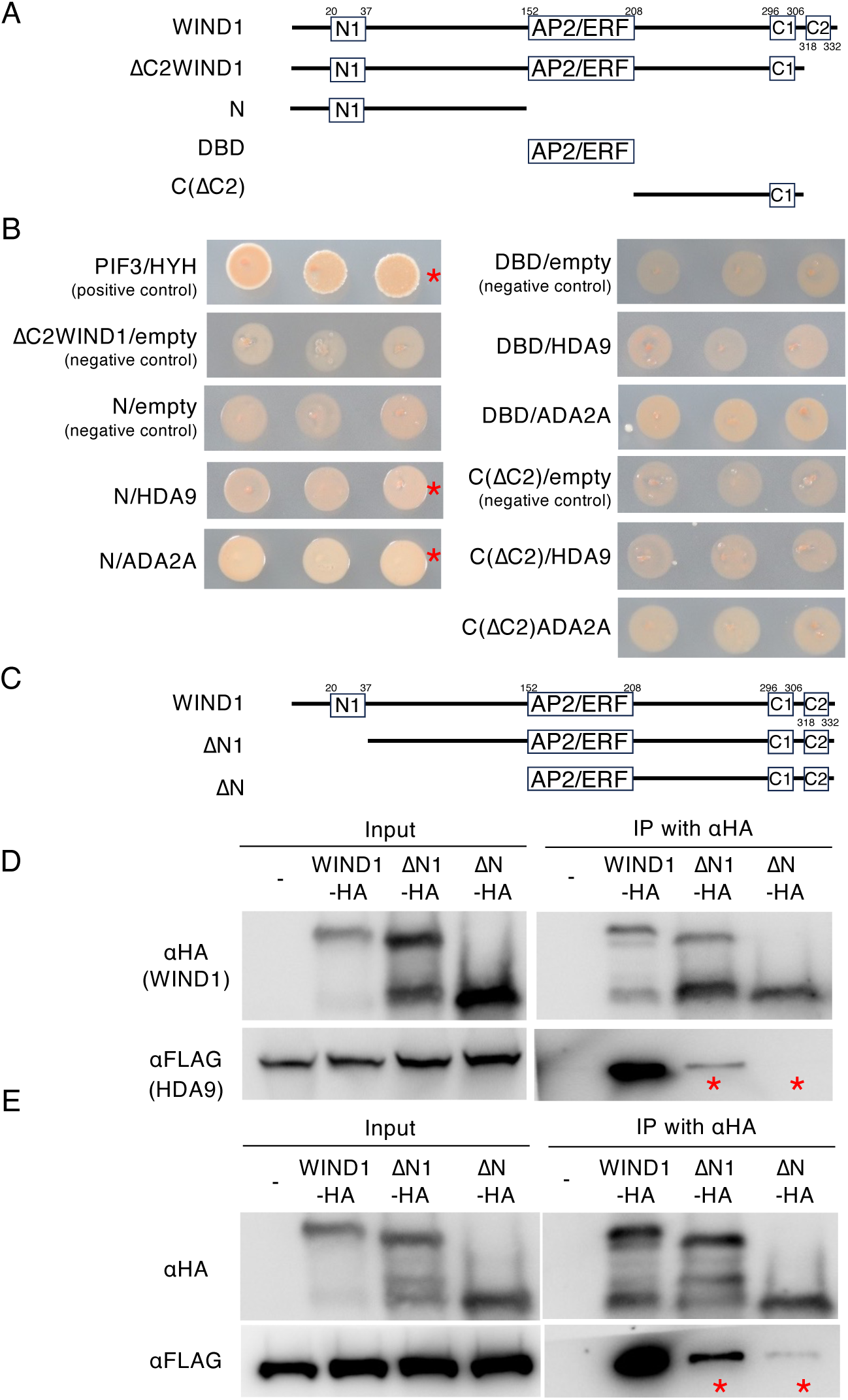
The N1 domain of WIND1 is responsible for interaction with HDA9 and ADA2a. (A) Schematic diagram of *WIND1* deletion series constructs. White boxes represent amino acid sequences conserved among WIND1 homologues (Iwase et al., 2013) and numbers above boxes show the order of amino acids. N1; conserved N-terminal domain 1, AP2/ERF; DNA binding domain, C1 and C2; conserved C-terminal domain 1 and 2, WIND1; WIND1 full-length, ΔC2WIND1; WIND1 lacking the C2 domain, N; WIND1 N-terminal region with 1^st^ to 151^st^ amino acids, DBD; WIND1 AP2/ERF DNA binding domain with 152^nd^ to 208^th^ amino acids, C(ΔC2); WIND1 C-terminal region with the 209^th^ to 317^th^ amino acids. (B) Y2H data showing the interaction between WIND1 N-terminal region and HDA9 or ADA2a. Transformed yeast cells grown on SD-Trp/-Leu/-Ade /-His/+AbA medium. PIF3-HYH binding was used as a positive control (Favero et al., 2020). The combination of ΔC2WIND1 and empty prey vector, N and empty prey vector, DBD and empty prey vector, and C(ΔC2) and empty prey vector served as negative controls. Asterisks indicate positive yeast growth. (C) Schematic diagram of WIND1 constructs. WIND1: full-length WIND1; ΔN1: WIND1 lacking amino acids 1–37; ΔN: WIND1 lacking amino acids 1–151 (D and E) Immunoprecipitation assay showing in vivo interactions of WIND1 with HDA9 (D) and ADA2a (E). Soluble extracts (input) from agroinfiltrated *N. benthamiana* leaves co-expressing WIND1-HA or its truncated versions (ΔN1-HA and ΔN-HA) with HDA9-FLAG (D) or ADA2a-FLAG (E) were subjected to immunoprecipitation with anti-HA antibody, followed by immunoblotting with anti-HA and anti-FLAG antibodies. Asterisks highlight reduced levels of HDA9-FLAG and ADA2a-FLAG co-immunoprecipitated with ΔN1-HA or ΔN-HA compared to full-length WIND1.

**Supplemental Figure 6.**
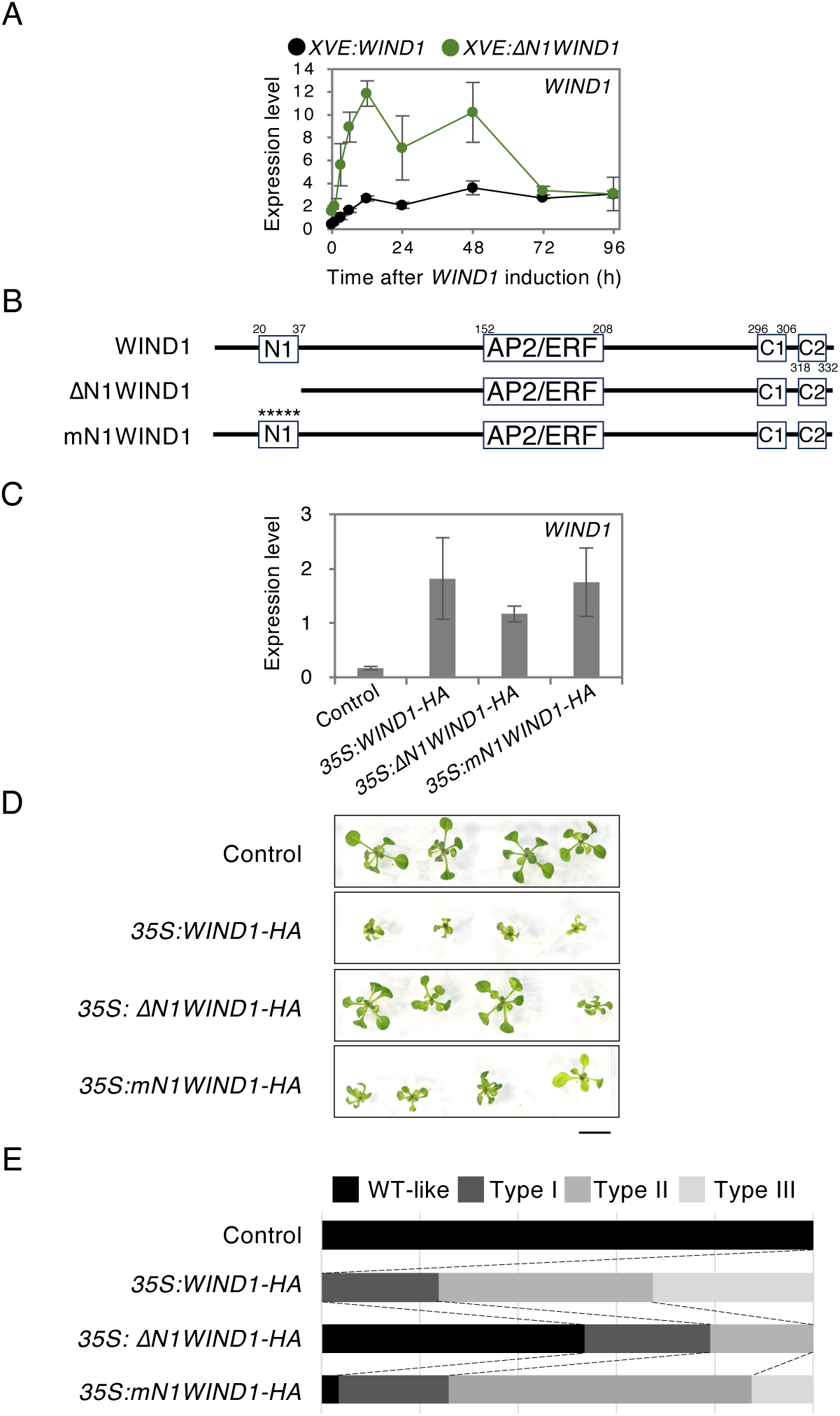
The N1 domain is required for WIND1-mediated cell fate reprogramming. (A) RT-qPCR data showing the transcript level of *WIND1* in *XVE:WIND1-HA* and *XVE:ΔN1WIND1-HA* plants used in Figure 2F and 2G. (B) Schematic diagram of *WIND1* constructs. White boxes represent amino acid sequences conserved among WIND1 homologues (Iwase et al., 2013) and numbers above boxes show the order of amino acids. N1; conserved N-terminal domain 1, AP2/ERF; DNA binding domain, C1 and C2; conserved C-terminal domain 1 and 2. WIND1 ; WIND1 full-length, ΔN1WIND1; WIND1 lacking from 1^st^ to the 37^th^ amino acids, mN1WIND1; WIND1 with amino acid substitution within the N1 domain. Asterisks highlight the substitution of two leucines (L21A, L25A) and three serines (S34A, S36A, S37A) with alanines (A) within the N1 domain. (C) RT-qPCR data showing the transcript level of *WIND1* in *35S:WIND1-HA*, *35S: ΔN1WIND1-HA* and *35S: mN1WIND1-HA* plants used in Supplemental Figure 6G and 6H. (D) Light micrograph of 21-day-old plants transformed with empty vector (Control), *35S:WIND1-HA*, *35S:ΔN1WIND1-HA* and *35S:mN1WIND1-HA* grown on phytohormone-free MS medium. Scale bar; 1 cm. (E) Quantitative data of morphologies alterations in (D). Phenotypic severities ranging from WT-like, weak (type I), intermediate (type II), and strong (type III) levels were scored according to Iwase et al. (2011).

**Supplemental Figure 7.**
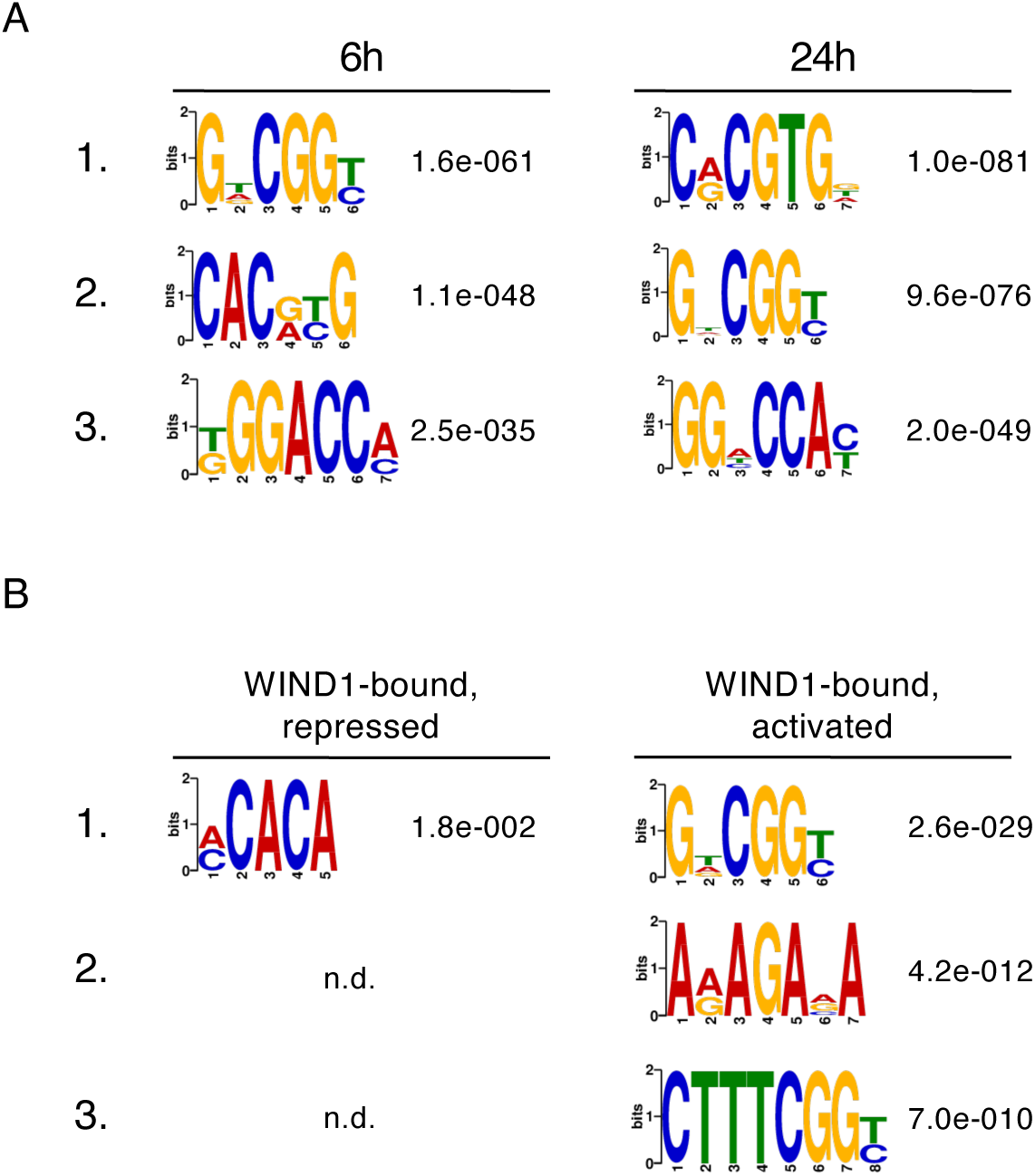
WIND1 preferentially binds to GC-rich G(T/A)CGG(T/C) and AC-rich (A/C)CACA motifs. (A and B) Sequence logos showing DNA motifs of 100 bp flanking sequences from WIND1-HA binding sites at 6 and 24 h after WIND1 induction (A). DNA motifs enriched in two distinct groups of WIND1 target genes defined by the Venn diagram in Figure 4A: genes that are directly bound by WIND1 and show decreased H3K27Ac and gene expression (114 genes, left), and genes that are directly bound by WIND1 and show increased H3K27Ac and gene expression (616 genes, right) (B). Top 3 motifs ordered by lower E-value, which is the enrichment *p*-value times the number of candidate motifs tested are shown. The enrichment *p*-value is calculated using Fisher’s Exact Test for enrichment of the motif in the positive sequences. n.d., not detected.

**Supplemental Table 1. A list of 2474 genes with decreased H3K27Ac levels at 3, 6, 24 and 48 h after *WIND1* induction in *XVE:WIND1* plants**

**Supplemental Table 2. A list of 4237 genes with increased H3K27Ac levels at 3, 6, 24 and 48 h after *WIND1* induction in *XVE:WIND1* plants**

**Supplemental Table 3. A list of 6615 genes downregulated at 1, 3, 6, 12, 24, 48, 72 and 96 h after *WIND1* induction in *XVE:WIND1* plants**

**Supplemental Table 4. A list of 4586 genes upregulated at 1, 3, 6, 12, 24, 48, 72 and 96 h after *WIND1* induction in *XVE:WIND1* plants**

**Supplemental Table 5. A list of 2069 genes that show WIND1-HA binding at 6 h after *WIND1* induction in *XVE:WIND1-HA* plants**

**Supplemental Table 6. A list of 2943 genes that show WIND1-HA binding at 24 h after *WIND1* induction in *XVE:WIND1-HA* plants**

**Supplemental Table 7. Oligonucleotides used in this study**

## REFERENCES

Adhikari, N. D., Froehlich, J. E., Strand, D. D., Buck, S. M., Kramer, D. M., and Larkin, R. M. (2011). GUN4-porphyrin complexes bind the chlH/GUN5 subunit of MG-chelatase and promote chlorophyll biosynthesis in Arabidopsis. Plant Cell 23:1449–1467.

Ali, M. A., Abbas, A., Kreil, D. P., and Bohlmann, H. (2013). Overexpression of the transcription factor RAP2.6 leads to enhanced callose deposition in syncytia and enhanced resistance against the beet cyst nematode Heterodera schachtii in Arabidopsis roots. BMC Plant Biol. 13:47.

Anzola, J. M., Sieberer, T., Ortbauer, M., Butt, H., Korbei, B., Weinhofer, I., Müllner, A. E., and Luschnig, C. (2010). Putative Arabidopsis transcriptional adaptor protein (PROPORZ1) is required to modulate histone acetylation in response to auxin. Proc. Natl. Acad. Sci. U. S. A. 107:10308–13.

Bannister, A. J., and Kouzarides, T. (2011). Regulation of chromatin by histone modifications. Cell Res. 21:381–395.

Birnbaum, K. D., and Alvarado, A. S. (2008). Slicing across Kingdoms: Regeneration in Plants and Animals. Cell 132:697–710.

Boutilier, K., Offringa, R., and Sharma, V. K. (2002). Ectopic expression of BABY BOOM triggers a conversion from vegetative to embryonic growth. Plant Cell 14:1737–1749.

Bouyer, D., Roudier, F., Heese, M., Andersen, E. D., Gey, D., Nowack, M. K., Goodrich, J., Renou, J.-P., Grini, P. E., Colot, V., et al. (2011). Polycomb repressive complex 2 controls the embryo-to-seedling phase transition. PLoS Genet. 7:e1002014.

Che, P., Lall, S., Nettleton, D., and Howell, S. (2006). Gene expression programs during shoot, root, and callus development in Arabidopsis tissue culture. Plant Physiol. 141:620–637.

Chen, Y., Hung, F. Y., and Sugimoto, K. (2023). Epigenomic reprogramming in plant regeneration: Locate before you modify. Curr. Opin. Plant Biol. 75:102415.

Chronis, C., Fiziev, P., Papp, B., Butz, S., Bonora, G., Sabri, S., Ernst, J., and Plath, K. (2017). Cooperative Binding of Transcription Factors Orchestrates Reprogramming. Cell 168:442–459.e20.

Clough, S. J., and Bent, A. F. (1998). Floral dip: a simplified method for Agrobacterium-mediated transformation of Arabidopsis thaliana. Plant J. 16:735–43.

Collins, C., Maruthi, N. M., and Jahn, C. E. (2015). CYCD3 D-type cyclins regulate cambial cell proliferation and secondary growth in Arabidopsis. J. Exp. Bot. 66:4595–4606.

Favero, D. S., Kawamura, A., Shibata, M., Takebayashi, A., Jung, J. H., Suzuki, T., Jaeger, K. E., Ishida, T., Iwase, A., Wigge, P. A., et al. (2020). AT-Hook Transcription Factors Restrict Petiole Growth by Antagonizing PIFs. Curr. Biol. 30:1454–1466.e6.

Feng, J., Liu, T., Qin, B., Zhang, Y., and Liu, X. S. (2012). Identifying ChIP-seq enrichment using MACS. Nat. Protoc. 7:1728–1740.

Gaj, M. D. (2001). Direct somatic embryogenesis as a rapid and efficient system for in vitro regeneration of Arabidopsis thaliana. Plant Cell Tissue Organ Cult. 64:39–46.

Heyman, J., Cools, T., Vandenbussche, F., Heyndrickx, K. S., Van Leene, J., Vercauteren, I., Vanderauwera, S., Vandepoele, K., De Jaeger, G., Van Der Straeten, D., et al. (2013). ERF115 controls root quiescent center cell division and stem cell replenishment. Science 342:860–3.

Heyman, J., Cools, T., Canher, B., Shavialenka, S., Traas, J., Vercauteren, I., Van Den Daele, H., Persiau, G., De Jaeger, G., Sugimoto, K., et al. (2016). The heterodimeric transcription factor complex ERF115-PAT1 grants regeneration competence. Nat. Plants 2:1–7.

Hung, F. Y., Feng, Y. R., Hsin, K. T., Shih, Y. H., Chang, C. H., Zhong, W., Lai, Y. C., Xu, Y., Yang, S., Sugimoto, K., et al. (2023). Arabidopsis histone H3 lysine 9 methyltransferases KYP/SUVH5/6 are involved in leaf development by interacting with AS1-AS2 to repress KNAT1 and KNAT2. *Commun*. Biol. 6:1–13.

Ikeda-Iwai, M., Umehara, M., Satoh, S., and Kamada, H. (2003). Stress-induced somatic embryogenesis in vegetative tissues of Arabidopsis thaliana. Plant J. 34:107–14.

Ikeuchi, M., Sugimoto, K., and Iwase, A. (2013). Plant Callus: Mechanisms of Induction and Repression. Plant Cell 25:3159–3173.

Ikeuchi, M., Iwase, A., Rymen, B., Harashima, H., Shibata, M., Ohnuma, M., Breuer, C., Morao, A. K., de Lucas, M., De Veylder, L., et al. (2015). PRC2 represses dedifferentiation of mature somatic cells in Arabidopsis. Nat. Plants 1:15089.

Ikeuchi, M., Ogawa, Y., Iwase, A., and Sugimoto, K. (2016). Plant regeneration: cellular origins and molecular mechanisms. Development 143:1442–1451.

Ikeuchi, M., Favero, D. S., Sakamoto, Y., Iwase, A., Coleman, D., Rymen, B., and Sugimoto, K. (2019). Molecular Mechanisms of Plant Regeneration. Annu. Rev. Plant Biol. 70:377–406.

Ikeuchi, M., Iwase, A., Ito, T., Tanaka, H., Favero, D. S., Kawamura, A., Sakamoto, S., Wakazaki, M., Tameshige, T., Fujii, H., et al. (2022). Wound-inducible WUSCHEL-RELATED HOMEOBOX 13 is required for callus growth and organ reconnection. Plant Physiol. 188:425–441.

Iwase, A., Mitsuda, N., Koyama, T., Hiratsu, K., Kojima, M., Arai, T., Inoue, Y., Seki, M., Sakakibara, H., Sugimoto, K., et al. (2011). The AP2/ERF transcription factor WIND1 controls cell dedifferentiation in arabidopsis. Curr. Biol. 21:508–514.

Iwase, A., Mitsuda, N., Ikeuchi, M., Ohnuma, M., Koizuka, C., Kawamoto, K., Imamura, J., Ezura, H., and Sugimoto, K. (2013). Arabidopsis WIND1 induces callus formation in rapeseed, tomato, and tobacco. Plant Signal. Behav. 8:1–5.

Iwase, A., Harashima, H., Ikeuchi, M., Rymen, B., Ohnuma, M., Komaki, S., Morohashi, K., Kurata, T., Nakata, M., Ohme-Takagi, M., et al. (2017). WIND1 Promotes Shoot Regeneration through Transcriptional Activation of ENHANCER OF SHOOT REGENERATION1 in Arabidopsis. Plant Cell 29:54–69.

Iwase, A., Kondo, Y., Laohavisit, A., Takebayashi, A., Ikeuchi, M., Matsuoka, K., Asahina, M., Mitsuda, N., Shirasu, K., Fukuda, H., et al. (2021). WIND transcription factors orchestrate wound-induced callus formation, vascular reconnection and defense response in Arabidopsis. New Phytol. 232:734–752.

Jopling, C., Boue, S., and Izpisua Belmonte, J. C. (2011). Dedifferentiation, transdifferentiation and reprogramming: three routes to regeneration. Nat. Rev. Mol. Cell Biol. 12:79–89.

Kadokura, S., Sugimoto, K., Tarr, P., Suzuki, T., and Matsunaga, S. (2018). Characterization of somatic embryogenesis initiated from the Arabidopsis shoot apex. Dev. Biol. Advance Access published 2018, doi:10.1016/j.ydbio.2018.04.023.

Kubo, M., Udagawa, M., Nishikubo, N., Horiguchi, G., Yamaguchi, M., Ito, J., Mimura, T., Fukuda, H., and Demura, T. (2005). Transcription switches for protoxylem and metaxylem vessel formation. Genes Dev. 19:1855–1860.

Langmead, B., and Salzberg, S. L. (2012). Fast gapped-read alignment with Bowtie 2. Nat. Methods 9:357–359.

Lee, K., Yoon, H., Park, O. S., Lim, J., Kim, S. G., and Seo, P. J. (2024). ESR2–HDA6 complex negatively regulates auxin biosynthesis to delay callus initiation in Arabidopsis leaf explants during tissue culture. Plant Commun. 5:100892.

Li, J., Zhang, Q., Wang, Z., and Liu, Q. (2024). The roles of epigenetic regulators in plant regeneration: Exploring patterns amidst complex conditions. Plant Physiol. 194:2022–2038.

Liu, X., Zhu, K., and Xiao, J. (2023). Recent advances in understanding of the epigenetic regulation of plant regeneration. aBIOTECH Advance Access published 2023, doi:10.1007/s42994-022-00093-2.

Lotan, T., Ohto, M., Yee, K. M., West, M. a, Lo, R., Kwong, R. W., Yamagishi, K., Fischer, R. L., Goldberg, R. B., and Harada, J. J. (1998). Arabidopsis LEAFY COTYLEDON1 is sufficient to induce embryo development in vegetative cells. Cell 93:1195–205.

Machanick, P., and Bailey, T. L. (2011). MEME-ChIP: Motif analysis of large DNA datasets. Bioinformatics 27:1696–1697.

Mahendrawada, L., Warfield, L., Donczew, R., and Hahn, S. (2025). Low overlap of transcription factor DNA binding and regulatory targets. Nature Advance Access published 2025, doi:10.1038/s41586-025-08916-0.

Mal, A., and Harter, M. L. (2003). MyoD is functionally linked to the silencing of a muscle-specific regulatory gene prior to skeletal myogenesis. Proc. Natl. Acad. Sci. U. S. A. 100:1735–1739.

Mao, Y., Pavangadkar, K. A., Thomashow, M. F., and Triezenberg, S. J. (2006). Physical and functional interactions of Arabidopsis ADA2 transcriptional coactivator proteins with the acetyltransferase GCN5 and with the cold-induced transcription factor CBF1. Biochim. Biophys. Acta - Gene Struct. Expr. 1759:69–79.

Mi, H., Ebert, D., Muruganujan, A., Mills, C., Albou, L. P., Mushayamaha, T., and Thomas, P. D. (2021). PANTHER version 16: A revised family classification, tree-based classification tool, enhancer regions and extensive API. Nucleic Acids Res. 49:D394–D403.

Mitsuda, N., Seki, M., Shinozaki, K., and Ohme-takagi, M. (2005). The NAC Transcription Factors NST1 and NST2 of Arabidopsis Regulate Secondary Wall Thickenings and Are Required for Anther Dehiscence. Plant Cell 17:2993–3006.

Mizukami, Y., and Fischer, R. L. (2000). Plant organ size control: AINTEGUMENTA regulates growth and cell numbers during organogenesis. Proc. Natl. Acad. Sci. U. S. A. 97:942–947.

Mori, S., Oya, S., Takahashi, M., Takashima, K., Inagaki, S., and Kakutani, T. (2023). Cotranscriptional demethylation induces global loss of H3K4me2 from active genes in Arabidopsis. EMBO J. 42.

Nishino, N., Jose, B., Shinta, R., Kato, T., Komatsu, Y., and Yoshida, M. (2004). Chlamydocin-hydroxamic acid analogues as histone deacetylase inhibitors. Bioorg. Med. Chem. 12:5777–5784.

Oberkofler, V., Pratx, L., and Bäurle, I. (2021). Epigenetic regulation of abiotic stress memory: maintaining the good things while they last. Curr. Opin. Plant Biol. 61.

Ohme-Takagi, M., and Shinshi, H. (1995). Ethylene-inducible DNA binding proteins that interact with an ethylene-responsive element. Plant Cell Online 7:173–182.

Ohta, M., Matsui, K., Hiratsu, K., Shinshi, H., and Ohme-Takagi, M. (2001). Repression domains of class II ERF transcriptional repressors share an essential motif for active repression. Plant Cell 13:1959–68.

Puri, P. L., and Sartorelli, V. (2000). Regulation of muscle regulatory factors by DNA-binding, interacting proteins, and post-transcriptional modifications. J. Cell. Physiol. 185:155–173.

Ramírez, F., Dündar, F., Diehl, S., Grüning, B. A., and Manke, T. (2014). DeepTools: A flexible platform for exploring deep-sequencing data. Nucleic Acids Res. 42:187– 191.

Robinson, M. D., McCarthy, D. J., and Smyth, G. K. (2009). edgeR: A Bioconductor package for differential expression analysis of digital gene expression data. Bioinformatics 26:139–140.

Robinson, J. T., Thorvaldsdóttir, H., Winckler, W., Guttman, M., Lander, E. S., Getz, G., and Mesirov, J. P. (2011). Integrative genomics viewer. Nat. Biotechnol. 29:24–26.

Rymen, B., Kawamura, A., Lambolez, A., Inagaki, S., Takebayashi, A., Iwase, A., Sakamoto, Y., Sako, K., Favero, D. S., Ikeuchi, M., et al. (2019). Histone acetylation orchestrates wound-induced transcriptional activation and cellular reprogramming in Arabidopsis. *Commun*. Biol. 2:404.

Sainsbury, F., Thuenemann, E. C., and Lomonossoff, G. P. (2009). pEAQ: versatile expression vectors for easy and quick transient expression of heterologous proteins in plants. Plant Biotechnol. J. 7:682–693.

Sakamoto, Y., Kawamura, A., Suzuki, T., Segami, S., Maeshima, M., Polyn, S., De Veylder, L., and Sugimoto, K. (2022). Transcriptional activation of auxin biosynthesis drives developmental reprogramming of differentiated cells. Plant Cell 34:4348–4365.

Sako, K., Kim, J.-M., Matsui, A., Nakamura, K., Tanaka, M., Kobayashi, M., Saito, K., Nishino, N., Kusano, M., Taji, T., et al. (2015). Ky-2, a Histone Deacetylase Inhibitor, Enhances High-Salinity Stress Tolerance in *Arabidopsis thaliana*. Plant Cell Physiol. 57:pcv199.

Sarojam, R., Sappl, P. G., Goldshmidt, A., Efroni, I., Floyd, S. K., Eshed, Y., and Bowmana, J. L. (2010). Differentiating Arabidopsis shoots from leaves by combined YABBY activities. Plant Cell 22:2113–2130.

Shibata, M., Favero, D. S., Takebayashi, R., Takebayashi, A., Kawamura, A., Rymen, B., Hosokawa, Y., and Sugimoto, K. (2022). Trihelix transcription factors GTL1 and DF1 prevent aberrant root hair formation in an excess nutrient condition. New Phytol. 235:1426–1441.

Stone, S. L., Kwong, L. W., Yee, K. M., Pelletier, J., Lepiniec, L., Fischer, R. L., Goldberg, R. B., and Harada, J. J. (2001). LEAFY COTYLEDON2 encodes a B3 domain transcription factor that induces embryo development. Proc. Natl. Acad. Sci. U. S. A. 98:11806–11.

Sugiyama, M. (2015). Historical review of research on plant cell dedifferentiation. J. Plant Res. 128:349–359.

Takahashi, K., and Yamanaka, S. (2006). Induction of pluripotent stem cells from mouse embryonic and adult fibroblast cultures by defined factors. Cell 126:663–76.

Tanaka, E. M., and Reddien, P. W. (2011). The Cellular Basis for Animal Regeneration. Dev. Cell 21:172–185.

Temman, H., Sakamoto, T., Ueda, M., Sugimoto, K., Migihashi, M., Yamamoto, K., Tsujimoto-Inui, Y., Sato, H., Shibuta, M. K., Nishino, N., et al. (2023). Histone deacetylation regulates de novo shoot regeneration. PNAS Next 2:1–12.

Ueda, M., Matsui, A., Tanaka, M., Nakamura, T., Abe, T., Sako, K., Sasaki, T., Kim, J.-M., Ito, A., Nishino, N., et al. (2017). The Distinct Roles of Class I and II RPD3-Like Histone Deacetylases in Salinity Stress Response. Plant Physiol. 175:1760– 1773.

Wang, F.-X., Shang, G.-D., Wu, L.-Y., Xu, G.-Z., Zhao, X.-Y., and Wang, J.-W. (2020). Chromatin Accessibility Dynamics and a Hierarchical Transcriptional Regulatory Network Structure for Plant Somatic Embryogenesis. Dev. Cell 54:1–16.

Wu, C. J., Yuan, D. Y., Liu, Z. Z., Xu, X., Wei, L., Cai, X. W., Su, Y. N., Li, L., Chen, S., and He, X. J. (2023). Conserved and plant-specific histone acetyltransferase complexes cooperate to regulate gene transcription and plant development. Nat. Plants 9:442–459.

Yang, W., Zhai, H., Wu, F., Deng, L., Chao, Y., Meng, X., Chen, Q., Liu, C., Bie, X., Sun, C., et al. (2024). Peptide REF1 is a local wound signal promoting plant regeneration. Cell 187:3024–3038.e14.

Yu, G., Wang, L. G., and He, Q. Y. (2015). ChIP seeker: An R/Bioconductor package for ChIP peak annotation, comparison and visualization. Bioinformatics 31:2382– 2383.

Zhang, T.-Q., Lian, H., Tang, H., Dolezal, K., Zhou, C.-M., Yu, S., Chen, J.-H., Chen, Q., Liu, H., Ljung, K., et al. (2015). An Intrinsic MicroRNA Timer Regulates Progressive Decline in Shoot Regenerative Capacity in Plants. Plant Cell Online 27:tpc.114.135186.

Zhou, S., Jiang, W., Long, F., Cheng, S., Yang, W., Zhao, Y., and Zhoua, D. X. (2017). Rice homeodomain protein WOX11 recruits a histone acetyltransferase complex to establish programs of cell proliferation of crown root meristem. Plant Cell 29:1088– 1104.

Zuo, J., Niu, Q.-W., and Chua, N.-H. (2000). An estrogen receptor-based transactivator XVE mediates highly inducible gene expression in transgenic plants. Plant J. 24:265–273.

Zuo, J., Niu, Q. W., Frugis, G., and Chua, N. H. (2002). The WUSCHEL gene promotes vegetative-to-embryonic transition in Arabidopsis. Plant J. 30:349–359.

